# In vivo selection and glymphatic delivery of AAV5 capsids engineered to target human glial progenitor cells

**DOI:** 10.1101/2025.07.03.662612

**Authors:** Alexander Cona, Evan Newbold, Deniz Kesmen, Rajiv Snape, Jessica Danner, Nicholas White, William Borden, Abigail Iseson, Steven J. Schanz, Devin Chandler-Militello, Xiaojie Li, Jose C. Cano, John N. Mariani, Maiken Nedergaard, Abdellatif Benraiss, Steven A. Goldman

## Abstract

To establish a means of efficiently transducing human glial progenitor cells (hGPCs) in vivo with therapeutic transgenes, we targeted PDGFRA-driven Cre-recombinase expressing hGPCs in human glial chimeric mice with a library of capsid-modified, recombination-reported adeno-associated viruses (AAVs). PCR screening for gliotropic viral capsid sequences, filtered against visceral organs, identified a set of AAV5-based vectors that preferentially infected human GPCs and/or their derived astrocytes and oligodendrocytes in vivo, with minimal systemic infection. To maximize the intracerebral distribution of these viruses while minimizing their dosing and extracerebral spread, we paired their intracisternal delivery with systemic hypertonicity. This method exploited intracerebral glymphatic flow to bypass the blood-brain barrier, delivering AAV directly into the brain parenchyma. Glymphatic delivery of capsid-modified AAV5s, evolved on human GPCs in vivo, thus enables efficient, brain-wide transgene delivery to human glia and their progenitors in the adult brain, with minimal off-target transduction.

## INTRODUCTION

A broad variety of neurological disorders are characterized by atrophy and involution of the cerebral white matter, with oligodendrocytic loss and demyelination unaccompanied by adaptive or compensatory remyelination. In a number of these conditions – including disorders as varied as progressive multiple sclerosis (PMS), Huntington disease, juvenile onset schizophrenia and the periventricular leukomalacia of cerebral palsy – dysmyelination evolves despite the persistence of a pool of parenchymal glial progenitor cells (GPCs) ^1,2^. These glial progenitors – which are typically referred to as oligodendrocyte progenitor cells, but which in humans may give rise to astrocytes as well as oligodendrocytes – appear to lose mitotic and differentiation competence in the setting of these disorders, whether by virtue of mitotic exhaustion, as in PMS and late aging, or due to cell-intrinsic defects in maturational pathways, as in HD and schizophrenia ^3–5^. In order to target these cells for therapeutic manipulation, vectors able to deliver genetic payloads to these cells, whether specifically so or to neurons as well, and to target the CNS only, with little or no systemic transduction, are needed.

Adeno-associated viruses (AAVs) enable stable expression of therapeutic transgenes in cell types of interest, but natural AAV serotypes can infect many different cell types, leading to both nonspecific infection and transgene expression by undesired cellular targets. While transgene expression may be restricted by cell type-specific regulatory elements, the range of host cell infection by AAVs - while serotype-specific – is nonetheless typically far broader than specific phenotypes of interest. The resultant promiscuity in infection leads to undesired off-target effects and toxicities, which have impeded the clinical development of therapeutic AAV vectors.

To address this issue, a number of strategies have been developed for evolving serotype binding domains to target prospectively-defined cell types of interest. In particular, previous studies have developed approaches towards capsid evolution based on Cre recombinase-dependent identification of viruses evolved to target cells of interest ^6^. These studies have included multiplexed assessment of recombination-dependent viral variants with randomly mutagenized VP1 capsid sequences, allowing high throughput screening of variants for their relative infectivity of given cell types of interest ^7^. However, most of these studies have evolved viral variants either in murine transgenic reporters with phenotype-selective Cre expression in vivo, or on human cells in vitro; none have been specifically evolved against human cells in vivo. Yet the surface epitopes presented by human cells are quite distinct from those presented by mice, and epitopes expressed in vivo may differ substantially from those targeted in vitro. Moreover, while the brain has been an especially attractive target for such studies, most investigators have focused on developing AAVs able to target neuronal populations; only a few studies have focused on targeting glial cells. In that regard, one AAV vector that preferentially targets striatal oligodendrocytes has been developed in rats that also infects these cells in macaques ^8,9^, though this vector requires intracerebral administration, so that its extrastriatal phenotypic selectivity remains unclear, as does its human infectivity. More broadly, to our knowledge no prior studies have focused on human glia in vivo, and no vectors have yet been reported that efficiently target glial progenitor cells (GPCs), the precursor to both oligodendrocytes and astrocytes, which comprise 3-4% of all brain cells in adult humans and mice alike ^10–14^. To thus develop vectors targeting human GPCs and their progeny in vivo, we adapted the previously described M-CREATE selection strategy of Deverman, Gradinaru and colleagues ^6,7^, by which Cre recombinase-expressing cells are used to screen recombination-reported viral variants for cell tropism, but we did so by capsid evolution on Cre-expressing human GPCs in vivo. In particular, we screened viral candidates in human glial chimeric mice, that had been neonatally engrafted with human embryonic stem cell-derived GPCs expressing Cre under the control of PDGFRA, which in the brain is selectively expressed by hGPCs. Our viral candidates were engineered to include lox66/lox71-flanked sequences, whose Cre-dependent recombination in targeted Cre-expressing host cells allowed the PCR-based identification of those viral variants that had most efficiently infected hGPCs (**Figure 1**).

**Figure 1.**
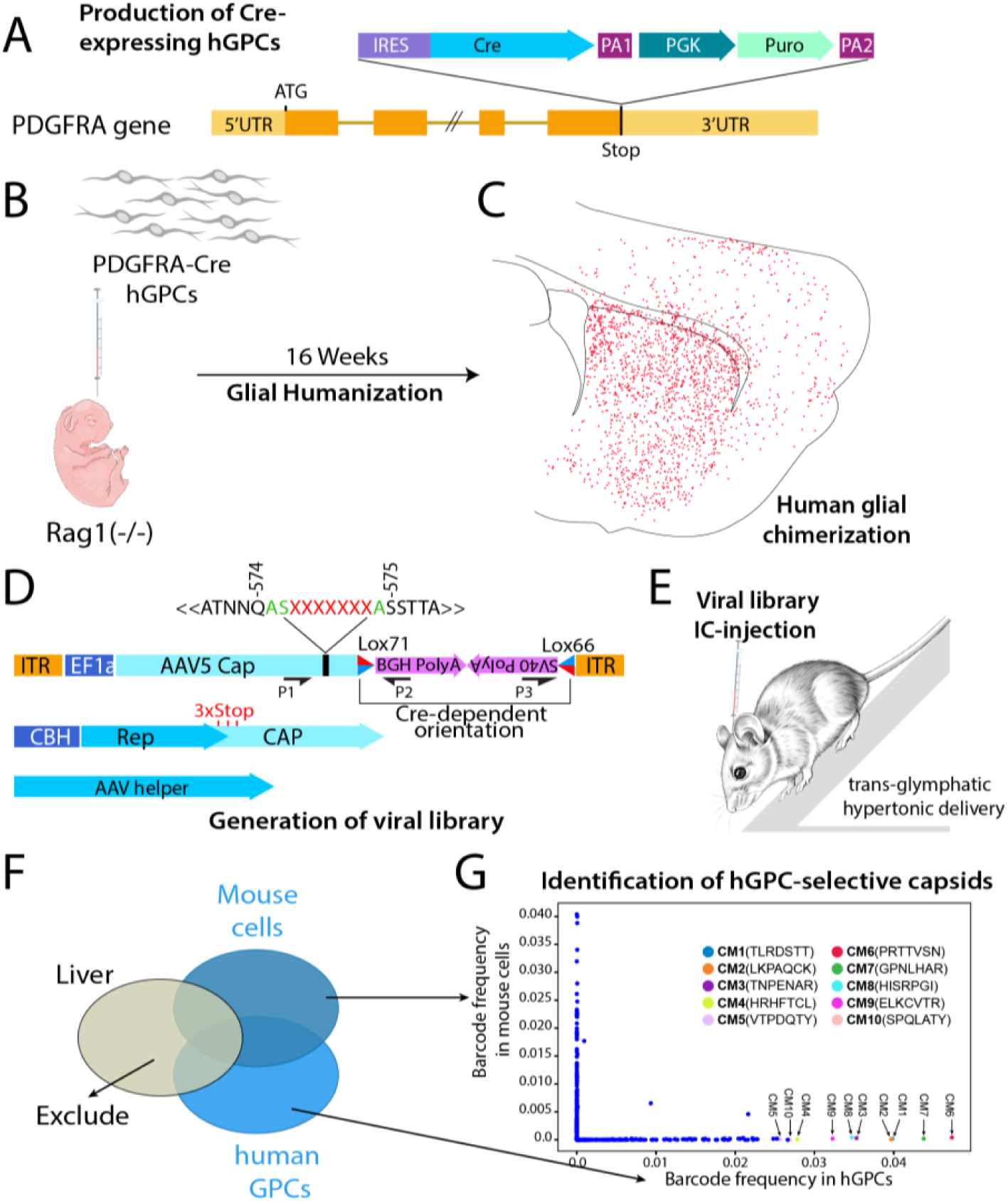
Production of capsid-evolved AAV5s designed to target human GPCs in vivo. **A.** hESCs were engineered to express PDGFRA-driven Cre by knocking in Cre 3’ to the endogenous PDGFRA locus. **B.** After glial induction, the human GPCs were injected into newborn rag1^−/−^ mice, to produce PDGFRA-Cre glial chimeras (**C**). **D** maps our approach towards generating a Cre-dependent AAV5 viral capsid library. A 21 base (7 mer) random sequence is placed between Q574/S575 of the VP1 capsid gene, in one of the VP1 loop binding domains. A Cre-sensitive Lox71/Lox66 flanked sequence harbors PCR primer sequences (P1-3) that are amplified, or not, depending upon whether Cre-triggered recombination has occurred in the infected human target cell. **E**, At 16 wks of age, the viral library was injected intracisternally into the brains of these mice (1 µl; 10^12^ vg/ml), under hypertonic conditions to potentiate glymphatic transport of virus. **F,** The mice were killed a week later, and brain and liver analyzed by PCR with primers for either Cre recombined (human GPCs; P1>3) or non-recombined sequences (mouse; P1>2). Capsid sequences recovered from liver were excluded from further analysis. **G**. Viral capsids in infected GPCs as a function of species; sequencing of PCR products showed that a number of viral variants were relatively specific to human GPCs, while others infected both mouse and human GPCs.

Yet the engineering of capsid-evolved vectors addresses only one aspect of the larger challenge of vector delivery to the brain; viral vectors, however efficient and cell type-selective, are only as good as their tissue access. AAV delivery to the brain has typically been limited by poor blood-brain barrier permeability and broad systemic leakage, requiring high – and hence toxic – doses. The blood-brain barrier in particular poses a formidable obstacle to the efficient transduction of neurons and glia by systemically-administered viral vectors, as it hinders viral entry from the systemic circulation. While serotypes such as AAV9-PHP.B and capsid-modified variants have been engineered to traverse the barrier^15–17^, their clinical use has nonetheless been limited by off-target transduction and the attendant toxicity of systemic infection ^18^, as well as by the immunotoxicity associated with high dose intravenous administration^19–23^. Thus, despite the development of a number of creative strategies towards circumventing the BBB, the targeting efficiencies for desired phenotypes remain relatively low, and to our knowledge, no vector has yet been reported to accomplish both BBB permeance *and* cell type-selective glial infection.

To establish a practicable strategy for circumventing the BBB using cell type-selective vectors, we therefore developed a method to co-opt the brain’s glymphatic system^24^ for viral delivery. By combining intracisternal injection with systemic hyperosmolarity^25,26^, we achieved broad AAV distribution throughout the brain, while avoiding BBB impedance. This delivery platform enabled the in vivo screening of a capsid library targeting hGPCs in human glial chimeric mice. We then filtered candidate vectors to exclude those with significant off-target hepatic infection, to identify variants that selectively and efficiently transduced hGPCs and their astrocytic and oligodendrocytic progeny. This approach yielded a panel of gliotropic viral vectors, each optimized to preferentially target a specific state along the human macroglial lineage. Combined with glymphatic delivery, these vectors demonstrate high on-target efficiency in vivo and with minimal systemic infection, offering a powerful platform for gene delivery to human glial cells in vivo.

## RESULTS

### Human glial chimeric mice may be established with Cre-expressing human GPCs

To establish human glial chimeric mice in which human GPCs could be identified as such following cell-specific CRE-dependent recombination, we first engineered human embryonic stem cells (hESCs) to express Cre recombinase under the control of PDGFRA, which is selectively expressed by glial progenitor cells in the adult brain ^27^. To this end, the CRE gene was knocked into the endogenous PDGFRA locus of hESCs (Genea19 line) by homology directed repair (HDR)-mediated insertion of donor DNA^28^. After sequence confirmation of the insert, and confirmation by array CGH that no major structural variants had been introduced into the Cre-expressing hESC line during editing, the cells were then differentiated into hGPCs using our described protocols ^29^. At 140-160 DIV, the hGPCs were transplanted (2 × 10^5^ cells/brain, in 4 divided injections of 5 × 10^4^ cells each) into the forebrains of neonatal Rag1^−/−^ immunodeficient mice; this technique allows the large-scale replacement of mouse by human GPCs, yielding chimeras in which most GPCs, and large numbers of their derived astrocytes, are human ^29–31^, while none differentiate as neurons (**Extended Data Figures 1A-D**). We generated 6 human glial chimeric mice for this set of experiments, that were colonized with PDGFRA-expressing hGPCs expressing Cre recombinase. Histological assessment of the hGPC-engrafted mice at 14 weeks of age confirmed widespread engraftment by human donor cells, and fluorescence in situ hybridization using RNAScope confirmed the expression of Cre recombinase mRNA by these cells (**Extended Data Figures 1E-H**).

### Hypertonic intracisternal delivery of AAV5 achieves widespread parenchymal access

We next asked whether we might use glymphatic pathways to access perivascular routes on the brain side of the BBB, so as to efficiently deliver AAV5 directly to the brain parenchyma. The glymphatic system comprises the network of perivascular channels by which CSF enters and pervades the brain, and by which this fluid exchanges with brain interstitial fluid to clear metabolic and protein wastes from the brain parenchyma^24,32^. Previous work had shown that glymphatic influx pathways may be used for compound delivery, just as its efflux pathways mediate metabolite clearance ^25^. We therefore asked if the glymphatic network might also serve as a route for viral delivery, circumventing the blood-brain barrier to allow direct parenchymal entry. We first assessed the utility and perivascular flow patterns of viral delivery via injection into the cisterna magna, which permits direct access to the glymphatic system via anterograde CSF flow ^26^. Critically, we paired intracisternal viral injection with both the systemic delivery of hypertonic saline and head-down prone (Trendelenburg) positioning, which together greatly increase glymphatic influx ^25^. We first used an FITC-coupled dextran as a tracer by which to validate the efficacy of this delivery approach, and observed widespread tracer entry throughout the forebrain (**Extended Data Figure 2**).

We next evaluated the utility of this pathway for viral administration. We assessed forebrain infection following the intracisternal hypertonic delivery of both an unmodified AAV5-serotyped AAV2 ITR-based vector (AAV2/5^33^, hereafter referred to as AAV5), and a modified AAV5 into whose VP1 capsid protein we inserted the neurotropic capsid sequence of AAV-retrograde (AAVrg)^34^, yielding AAV5-retro. Using our injection protocol, we observed widespread parenchymal infection by each of these control viruses. AAV5-retro in particular infected high proportions of hippocampal pyramidal and corticofugal neurons following a single intracisternal injection (**Extended Data Figures 3A-C**), yet unmodified AAV5 similarly infected primarily neurons; both control vectors exhibited relatively sparse glial labeling (**Extended Data Figures 3D-F**). Together, these experiments indicated that intracisternal administration of AAV5 under hypertonic conditions enables its rapid, brain-wide perivascular spread, circumventing the blood-brain barrier to allow direct viral entry into the brain with widespread, efficient cellular transduction. They also confirm prior observations that both unmodified AAV5 and modified AAVs bearing the retro capsid insert are primarily neurotropic^35^.

### AAV5 capsids may be evolved to target human glial progenitor cells in vivo

Once we had established chimeric mice whose incorporated human glia expressed Cre recombinase, and had optimized a delivery strategy for delivering AAVs bearing recombination-sensitive sequences, we next injected the human glial chimeric mice intracisternally with a viral library of candidate AAV5 capsids (1 µl at 1 × 10^12^ vg/ml, hence 1 × 10^6^ vg/injection). The library was generated by the ligation of a 21 random nucleotide sequence inserted after Q574 of VP1, a recognition loop on the sialic acid receptor binding pocket, so as to randomly modify the binding specificity of the capsid, as has been described ^36^. We inserted a GCT codon (alanine) between Q574 and S575 of the AAV5 VP1 capsid, to introduce an NheI cut site for insertion of the heptamer-encoding sequences of the library; the engineered AAVs thus carried random cap sequences mediating their binding specificity. As such, each of these viral candidates expressed a random heptapeptide insert in the VP1 binding domain, thereby permuting its differential adhesion to target cells. We estimated the resultant library to encode 4 × 10^5^ of these capsid variants. Importantly, the resultant AAV Cap sequences were placed upstream to two polyA tails, BGH and SV40 poly-adenylation sequences inserted in opposite directions, both flanked by two different Lox sequences (Lox66 and Lox71); this allowed the viral DNA to be detected by PCR as either unperturbed, or as the product of Cre-dependent reorientation, using PCR primers within each polyA sequence (**Figure 1D**). In virally-infected PDGFRA-Cre expressing human GPCs, the recombination of these sites allowed for the specific amplification of those viruses that had successfully transduced the human cells, while viruses that infected mouse cells remained unchanged.

As noted, 10 µl of this viral capsid library (1 × 10¹² vg/ml) were then injected intra-cisternally into each of four 16-week-old mice, after inducing systemic hypertonicity with an intraperitoneal infusion of NaCl (1M, or 5.8%; 20 µl/g b.w.), and with each mouse in a prone Trendelenburg position as described above (**Figure 1E**). The mice were euthanized a week after viral delivery, and their forebrain viral sequences were assessed by PCR using two sets of primers: one corresponding to the rearranged virion DNA resulting from recombination in hGPCs, where PDGFRA-driven Cre was active at the time of infection, and a second set corresponding to the native viral DNA sequence, which remained unchanged in the infected mouse cells. Importantly, to exclude viral capsids with significant off-target systemic affinities, the liver was also harvested from the infected mice, gDNA was extracted, and those capsids that had infected the liver were separately assessed by PCR for each mouse. This step allowed us to define and exclude those capsid variant sequences that targeted liver as well as brain, regardless of their relative affinity for hGPCs. We thereby identified those capsids most selective for human brain cells, and for hGPCs and their derivatives in particular, while manifesting the least off-target systemic hepatic infection **(Figures 1F-G).** Choosing the top 5-most human glia-enriched capsids recovered from each mouse allowed us to identify a set of 20 different capsid variants, which we designated CM1-20, that efficiently infected GPCs, and exhibited differential infectivity of human GPCs relative to mouse (**Table 1**).

**Table 1.**
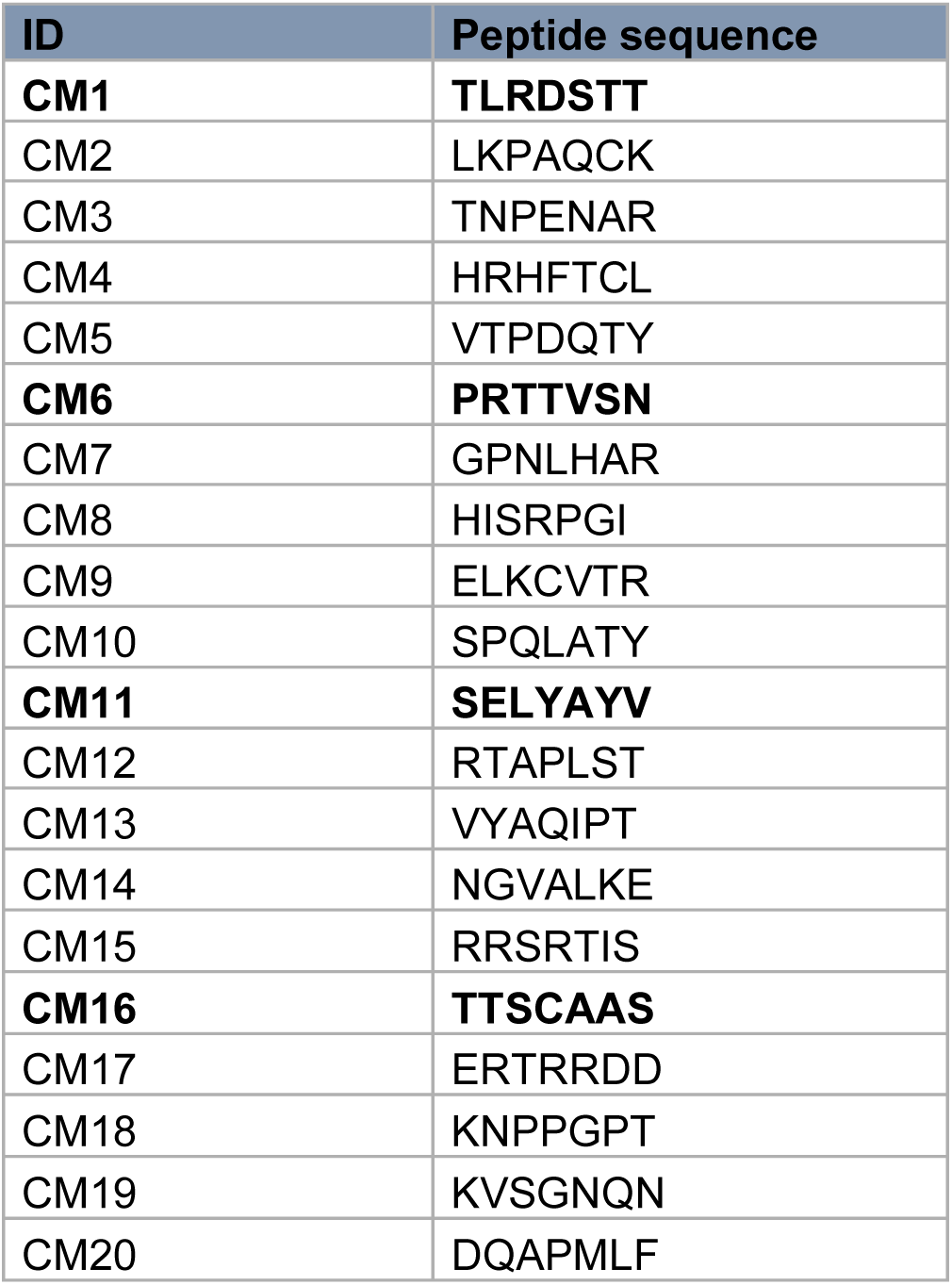
Capsid insert sequences. This table lists the peptide insertions in the AAV2/5 VP1 capsid that were most recovered from infected PDGFRA-Cre hGPCs after Cre-mediated recombination, comprising the 20 most human glial-selective peptide sequences that we recovered (see also **Figure 1**). CM1, 6, 11 and 16 (in **bold**) were the most human and glial selective among the recovered capsid inserts; among these, CM1 and CM6 were those that survived filtration against systemic organs, that exhibited the greatest human selectivity, and which infected most efficiently.

| ID | Peptide sequence |
| --- | --- |
| **CM1** | **TLRDSTT** |
| CM2 | LKPAQCK |
| CM3 | TNPENAR |
| CM4 | HRHFTCL |
| CM5 | VTPDQTY |
| **CM6** | **PRTTVSN** |
| CM7 | GPNLHAR |
| CM8 | HISRPGI |
| CM9 | ELKCVTR |
| CM10 | SPQLATY |
| **CM11** | **SELYAYV** |
| CM12 | RTAPLST |
| CM13 | VYAQIPT |
| CM14 | NGVALKE |
| CM15 | RRSRTIS |
| **CM16** | **TTSCAAS** |
| CM17 | ERTRRDD |
| CM18 | KNPPGPT |
| CM19 | KVSGNQN |
| CM20 | DQAPMLF |

### A family of capsid evolved AAV5s selectively infect GPCs and astrocytes

A number of glial ectodomains are co-expressed by astrocytes as well as their progenitors, and as such may also be recognized by viruses that infect PDGFαR-expressing GPCs. As a result, we found that our hGPC-selective clones manifested some degree of astrocytic targeting as well - some minimal, such as CM1 (**Figures 2A-E**), and some profoundly so, such as CM6, which selectively targeted both GPCs and astrocytes efficiently (**Figures 2F-G**). Focusing our efforts on these two vectors, after initial screening that identified them as transducing mouse as well as human glia, we found that while some neuronal transduction was exhibited by each, both demonstrated marked gliotropism relative to the unmodified AAV5s from which they were derived (**Figures 2I-K**). In contrast, in controls injected with unmodified WT AAV5, almost 50% of infected cells were neurons (**Extended Data Figure 4A**); thus, both AAV-CM1 and AAV-CM6 exhibited significantly more avidity for glia than did WT AAV5 (n=3-4 mice/group; p<0.0001, 1-way ANOVA with post hoc Tukey’s t-tests). That said, neuronal infection by these AAV5-derived vectors was quantitatively weak; in the same brains and regions in which glial infection was scored, less than 1% of all neurons were infected (**Extended Data Figure 4B**). Together, these data serve to highlight both the relative gliotropism and greater net glial infectivity of our modified vectors, relative to unmodified AAV5.

**Figure 2.**
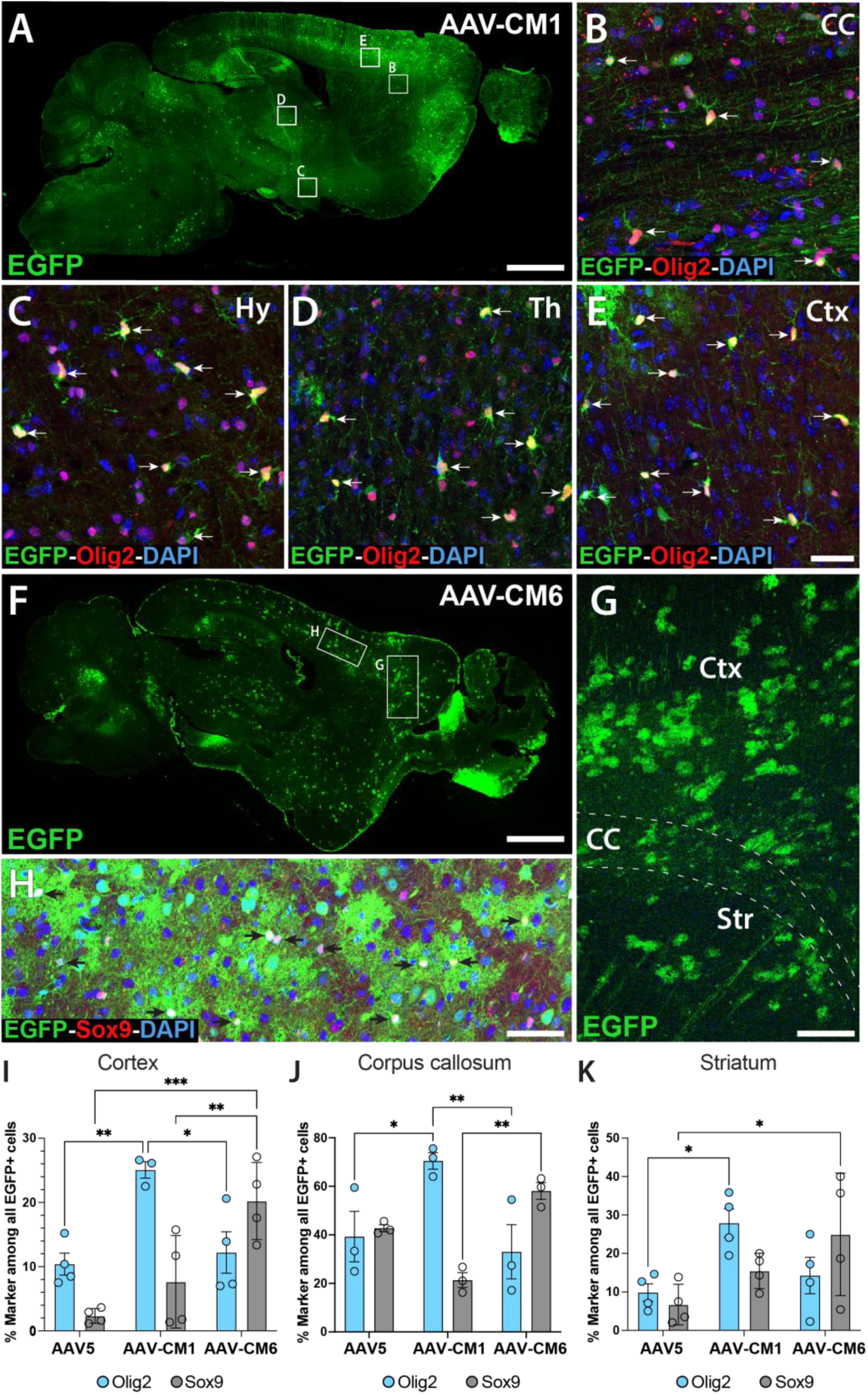
Different capsid variants exhibit distinct glial tropisms. Based on the sequence data of **Table 1**, AAV5s were engineered whose capsids presented two of the candidate peptides selected on PDGFRA-Cre expressing hGPCs in vivo. This figure shows sagittal images of brains infected by one of two different capsids, AAV CM1 (**A-E**) and CM6 (**F-H**). In each case, virus was injected (10 µl; 1×10^12^ vg/ml) into the cisterna magna of 12 wk-old mice, that were rendered hypertonic (3% NaCl IV infusion) 15 min prior to intracisternal injection, and killed 3 wks later. EGFP^+^ infected glia were recognized throughout the brain. **A,** Low power sagittal view of AAV CM1-infected, EGFP-reported infected cells, scattered throughout the brain. **B-E,** show white matter fields in the corpus callosum (**B**), hypothalamus (**C**), thalamus (**D**) and cortex (**D**). In each example, most infected cells co-labeled for Olig2, which is expressed by GPCs as well as oligodendrocytes; few infected non-glia were noted in this animal. **F,** Low power view of AAV CM6-infected cells. **G,** view from cortex through callosum to striatum, showing infected cells; most are GPCs, with a few astrocytes. **H,** Most AAV CM6-infected cells expressed Sox9, indicating astrocytic phenotype. **I-K**, In all 3 regions assessed, including cortex (**I**), corpus callosum (**J**) and striatum (**K**), AAV-CM1 preferentially infected Olig2-defined hGPCs and oligodendroglial lineage cells, while AAV-CM6 preferentially targeted Sox9-defined astrocytes. Two-way ANOVA with Sidak’s post-hoc tests. Means ± SEM. *p<0.05, **p<0.01, ***p<0.001. Scale, **A, F**: 1 mm. **B-E**, **H**: 50 µm. **G**: 200 µm.

### Capsid-evolved AAVs were filtered to exclude those manifesting systemic infection

While there is scant literature concerning gliotropic AAVs, a number of neurotropic AAVs that have advanced to trails - including those modified from AAV9 that cross the blood-brain barrier - have been associated with significant toxicity, largely due to adventitious infection of the liver and dorsal root ganglia, yielding hepatic dysfunction and pain syndromes respectively^18^. This may be the product of the high systemic doses need to achieve BBB passage and the large fraction of those doses that adventitiously infect off-target tissue, such as the liver and spleen, with consequent immune activation. Our use of glymphatic delivery to circumvent the BBB allows significantly lower viral doses, thus mitigating toxicity. Nonetheless, viral escape from the CNS may permit viral access to these same off-target tissues.

To minimize the possibility of non-CNS infection, we screened our vectors to identify those with the least infection of systemic tissues. To this end, we included PCR screening for viral sequence of the livers of each animal in which viral candidates were assessed, in light of that organ’s frequent off-target infection by AAVs and associated toxicity. The hepatic viral genome concentrations (vg/g) were compared to that of the targeted brains of the same animals, so as to assess the degree of off-target infection by each of our candidate capsid variants. All candidates manifesting appreciable hepatic infection were excluded from further analysis; this step served as a gatekeeper to identify those capsid variants appropriate for further development and in vivo assessment. We found that the capsid candidates that survived this filtration, delivered intracisternally, exhibited little or no infection of the liver, or of the spleen or kidneys either (**Extended Data Figures 5A-C**). Two of these vectors in particular, CM1 and CM6, appeared to allow phenotype-restricted delivery of reporter transgenes to brain glia in vivo, with minimal visceral infection. Together, these data indicate that hGPC-targeted capsid-modified AAV5s, in particular our prototypic vectors CM1 and CM6, exhibit substantially greater glial infectivity than wild-type AAV5, with lesser though persistent infectivity of neuronal populations, and little if any off-target visceral infection.

### Capsids evolved on human GPCs exhibited efficient human glial infectivity in vivo

The data described thus far describe the relative infectivities of our hGPC capsid-evolved vectors on mouse glia in non-chimeric mice; these experiments demonstrated the marked gliotropism of these vectors relative to the wild-type, unmodified AAV5 from which our variants were derived. We next asked if these vectors could efficiently target *human* GPCs and their derivatives in vivo, and if so, whether human cells were *selectively* infected relative to their murine counterparts. To that end, we compared the glial targeting efficacy of our capsid-modified AAV5-CM1 to that of wild-type (WT) unmodified AAV5 (AAV2/5), and did so in human glial cells in vivo. In particular, we engineered both WT AAV5 and AAV5-CM1 vectors to express distinct fluorescent reporters in a Cre-dependent manner, and then compared the relative infectivities of these viruses head-to-head in hGPCs in vivo, in PDGFRα-Cre human glial chimeric mice. For this purpose, the flex-EGFP reporter ^37^ was packaged into AAV-CM1, while the flex-dtTomato reporter was packaged into WT AAV5 (**Figure 3A**); the two reporter viruses were prepared in equal concentrations, and equal volumes then mixed together. A total of 10 µl (1 × 10^12^ vg/ml) of this 1:1 mixture was then injected into the cisterna magna of 12-month-old mice (n=3), each of which had been neonatally chimerized with PDGFRα-Cre expressing hGPCs. Three weeks later the mice were sacrificed, and the relative expression of each vector was assessed in the engrafted human cells (**Figures 3B-C**). This analysis revealed that AAV-CM1-EGFP transduced a significantly and substantially higher proportion of hGPCs than did AAV5-dtTomato (**Figure 3D**), confirming the superior glial avidity of the AAV-CM1 vector for human glia in vivo.

**Figure 3.**
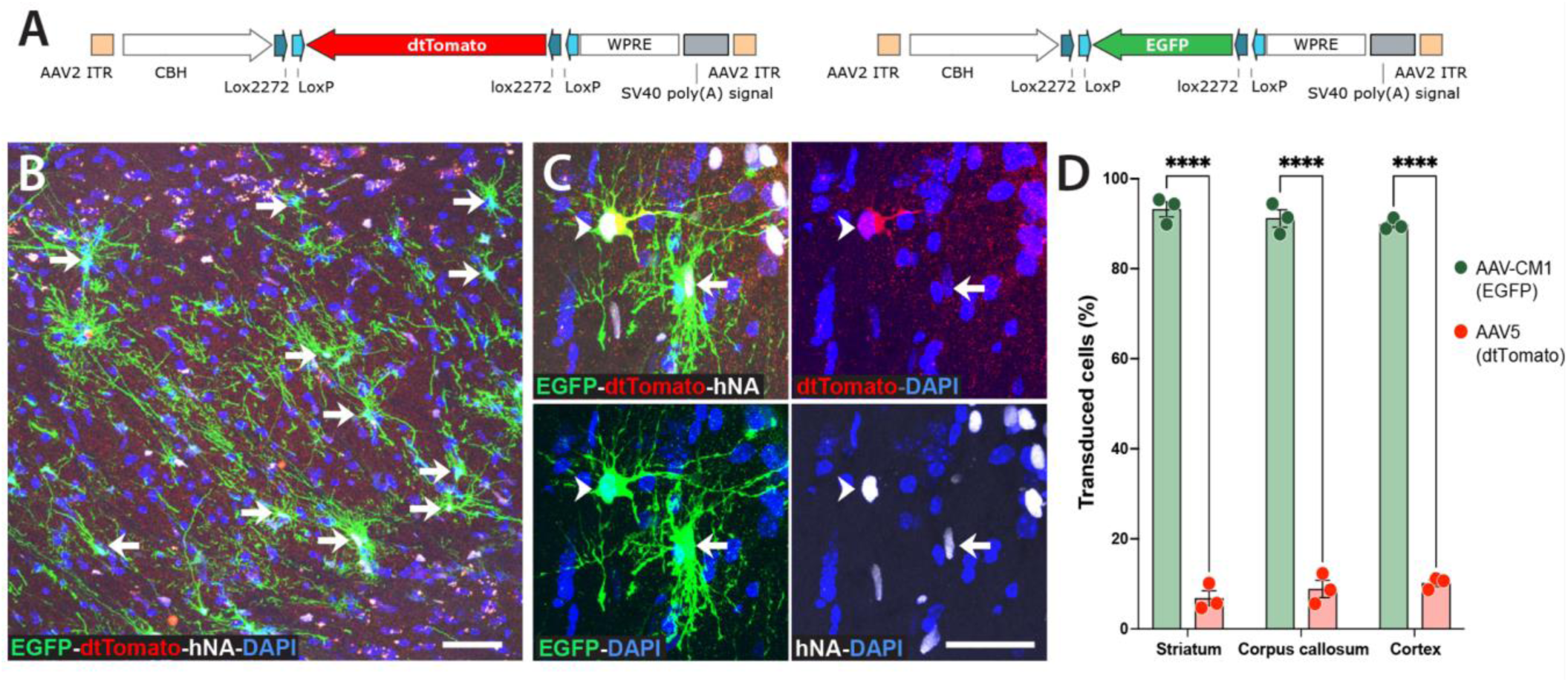
Viral selection on human PDGFRA-Cre GPCs in vivo yields vectors selective for glia. AAV-CM1 was significantly more efficient than its parental AAV5 (AAV2/5, an AAV2 ITR-based vector pseudotyped with a wild-type AAV5 capsid) in transducing hGPCs in vivo. **A.** Schematic of the viral constructs used to generate AAV-CM1 and AAV2/5 viruses. **B-C.** PDGFRα-Cre chimeric Rag1^−/−^ mice (transplanted with PDGFRA-Cre hGPCs on postnatal day 1) were injected at 12 months of age with a 10 µl viral cocktail containing AAV5 (AAV2/5 WT)-Flex-dtTomato and AAV-CM1-Flex-EGFP (1:1 ratio; 1 × 10^12^ vg/ml). The mice were sacrificed three weeks after injection. Brain sections were immunostained with anti-EGFP, anti-mCherry (recognizing dtTomato), and anti-human nuclear antigen (hNA) antibodies. **D.** Quantification of cells expressing EGFP and dtTomato in graft-derived hGPCs from the striatum, corpus callosum, and cortex. Means ± SEM. Two-way ANOVA with Sidak’s post-hoc test (****p <0.0001). Scale, 50 µm.

Having established the enhanced tropism of AAV-CM1 for human glia in vivo, relative to wild-type AAV, we next assessed the respective phenotypic biases of our most efficient vectors, CM1 and CM6. We again used adult human glial chimeric mice, that had been perinatally engrafted with hGPCs produced from unmodified hESCs (line WA09), to assess the avidity of these vectors; we did so by injecting the viruses intracisternally at 16 weeks of age with either AAV-CM1 or AAV-CM6, and sacrificing them 3 weeks later to assess both their transduction efficiencies and preferred phenotypic targets. We found that CM1 exhibited significant bias towards hGPC and oligodendroglial lineage infection, while CM6 was especially tropic for astrocytes (**Figures 4A-C; 4E-G)**. In the corpus callosum, AAV-CM1 transduced 5.3-fold more human Olig2^+^-defined glia than did the parental WT AAV5: 34.9% ± 5.7 of all human Olig2^+^ glia in the scored field were infected by a single injection of AAV-CM1, whereas only 6.5% ± 1.1 of these cells were infected in WT AAV5-injected mice (n=4/group; means ± SEM, p=0.0014, 2-way ANOVA with Sidak’s multiple comparison test) (**Figure 4D**). In contrast, AAV-CM6 transduced 3.8-fold more Sox9^+^-defined astroglia than did WT AAV5 (AAV-CM6: 22.2% ± 3.9; AAV5: 5.9% ± 2.6; n=4/group, means ± SEM, p=0.002, 2-way ANOVA) (**Figure 4H**). Thus, while neither CM1 nor CM6 was exclusive for its preferred target, their respective very strong biases towards hGPCs and astrocytes were each highly significant.

**Figure 4.**
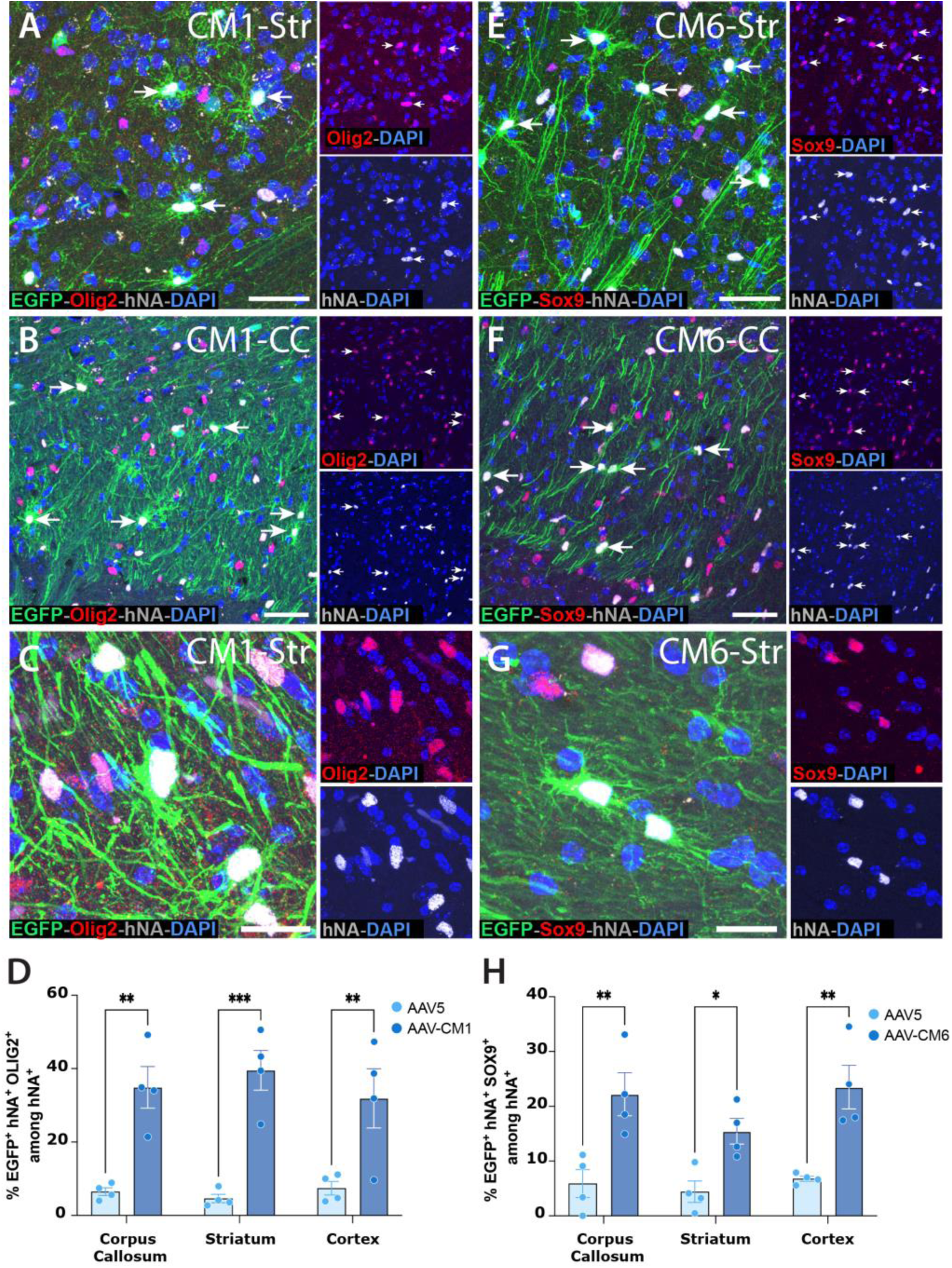
Capsid-evolved AAV5s CM1 and CM6 transduce distinct human glial phenotypes in vivo. Human glial chimeric mice, given neonatal intrastriatal injections of hESC-derived hGPCs, were injected intracisternally with AAV-CM1-CAG-EGFP or AAV-CM6-CAG-EGFP at 16 weeks of age, then examined 3 weeks later. EGFP expression in AAV-CM1-injected mice preferentially colocalized with Olig2 (**A-C**), whereas AAV-CM6 primarily infected Sox9-defined astrocytes (**E-G**). **D-H** show that AAV-CM1 and AAV-CM6 were significantly more efficient at targeting human glial progenitors and oligodendroglia (**D**) and astrocytes (**H**) than the unmodified AAV5 control vector. Two-way ANOVA with Sidak post-hoc tests. Means ± SEM. *p<0.05, **p<0.01, ***p<0.001. Scale: **A-B, E-F**:100 µm, **C, G**: 25 µm.

The phenotypic selectivity of these vectors was all the more remarkable in that each exhibited greater tropism to human than mouse cells of each phenotype (**Figure 5**). To assess the human-selectivity of these vectors, over and above their glial tropism, we assessed the relative infection of mouse and human GPCs in the corpus callosa of chimeric *shiverer* mice (*MBP^shi/shi^ x rag2^−/−^*) that were neonatally engrafted, injected with AAV-CM1 at 16-17 weeks, and killed 3 weeks later. In these immunodeficient and myelin-deficient mice, hGPCs competitively dominate the host mouse GPCs^38^, such that just over half (52.2 ± 6.0%, n=3) of all Olig2-defined hGPCs and oligodendroglia were of human origin at the time of sacrifice, consistent with our past studies of hGPC colonization of the neonatally-engrafted *shiverer* brain^39^ (**Figures 5A-D**). We found that of all Olig2-expressing cells in the scored corpus callosa, 87 ± 3.3% of the infected cells were human, and 13 ± 3% murine (n=3 mice/group; <0.0001 by t-test; **Figure 5D**). After normalization for the incidence of each species’ GPCs in the chimeric white matter, we found that of all human Olig2-expressing cells in the scored corpus callosa, 30.2 ± 1.1% were infected by CM1. In contrast, just 5.0 ± 0.2% of Olig2+ mouse cells were infected (n=3 mice/group; p<0.001 by t-test) (**Figure 5E**). As such, when co-resident in the same regions of corpus callosum in roughly the same incidence, the human GPCs were roughly 6-fold more likely to be infected by AAV CM1 than were mouse cells. This hGPC-evolved vector thus manifested strong selective tropism for human GPCs relative to mouse GPCs.

**Figure 5.**
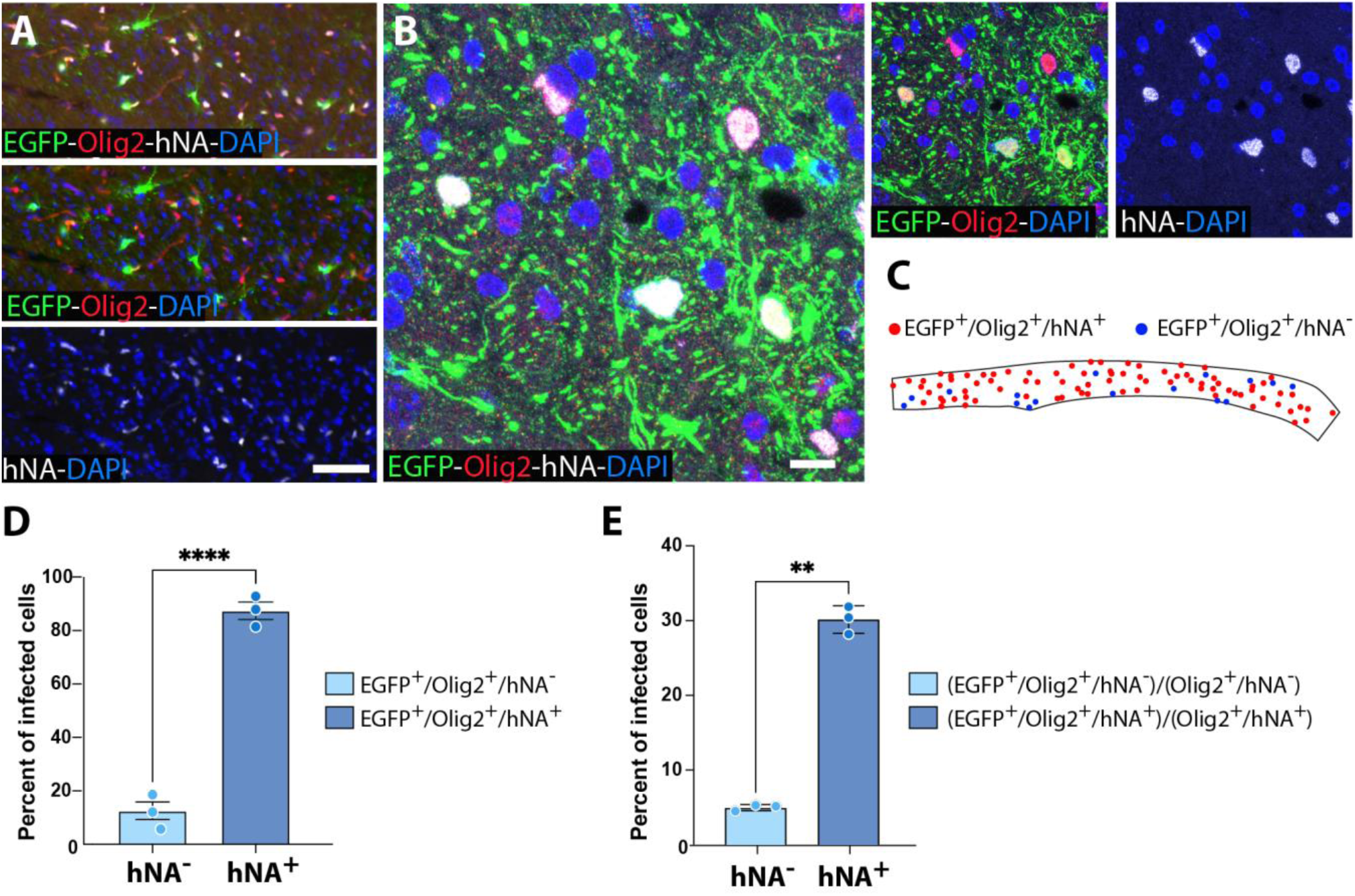
AAV-CM1 selectively transduces human glia. Neonatally chimerized shiverer mice, with hESC-derived glia engrafted into the striatum, were injected with AAV-CM1-EGFP virus into the cisterna magna at 17 weeks. Upon sacrifice and histological analysis three weeks later, we demonstrated that the virus preferentially transduces graft-derived human cells in the corpus callosum (**A**). Immunostaining for human nuclear antigen (hNA, white), Olig2 (red), and EGFP (green), as well as a DAPI (blue) counterstain. **B** and its color-split insets show a high-power field revealing co-localization of EGFP and Olig2 in human (hNA^+^) cells. **C.** Dot-mapped distribution in the corpus callosum of infected (EGFP^+^) human (hNA) and mouse Olig2^+^ cells. **D**. Graph plots the preferential infection of human relative to mouse glia. **E.** Plots the respective infection of human and mouse glia after normalization for the relative proportions of all human and mouse cells in the chimeric corpus callosa. Means ± SEM. **p <0.01 ****p <0.0001, two-tailed t-test. Scale bar: **A**: 10 µm; **B**: 100 µm.

### CM1 exhibited marked infection of mature oligodendrocytes as well as of their progenitors

In light of the efficient infection by AAV5-CM1 of hGPCs and Olig^+^ oligodendroglial lineage cells broadly defined, we next asked if this vector could efficiently infect mature oligodendrocytes, an especially important clinical target for which few competent delivery vectors have been described, and then only in rodent models. To this end, we established mice with largely humanized white matter, by transplanting hGPCs into neonatal immunodeficient shiverer mice (*MBP^shi/shi^ x rag2^−/−^*); this model permits the robust oligodendrocytic maturation and myelination by their engrafted hGPCs^29,30^. When the mice reached 16-17 weeks of age, by which time they typically exhibit a mature, largely humanized forebrain white matter, AAV5-CM1 was delivered intracisternally, after induction of systemic hypertonicity. When sacrificed 3 weeks later, robust infection of myelinated oligodendrocytes – as well as parenchymal astrocytes and persistent hGPCs - was noted in the corpus callosum and major forebrain white matter tracts (**Figure 6**). These data indicate the potential for therapeutic transgene delivery via AAV5-CM1 to mature myelinated oligodendroglia as well as to their progenitors, thus enabling a strategy for widespread transduction of human oligodendroglia in vivo, via a readily applicable route of intracisternal delivery.

**Figure 6.**
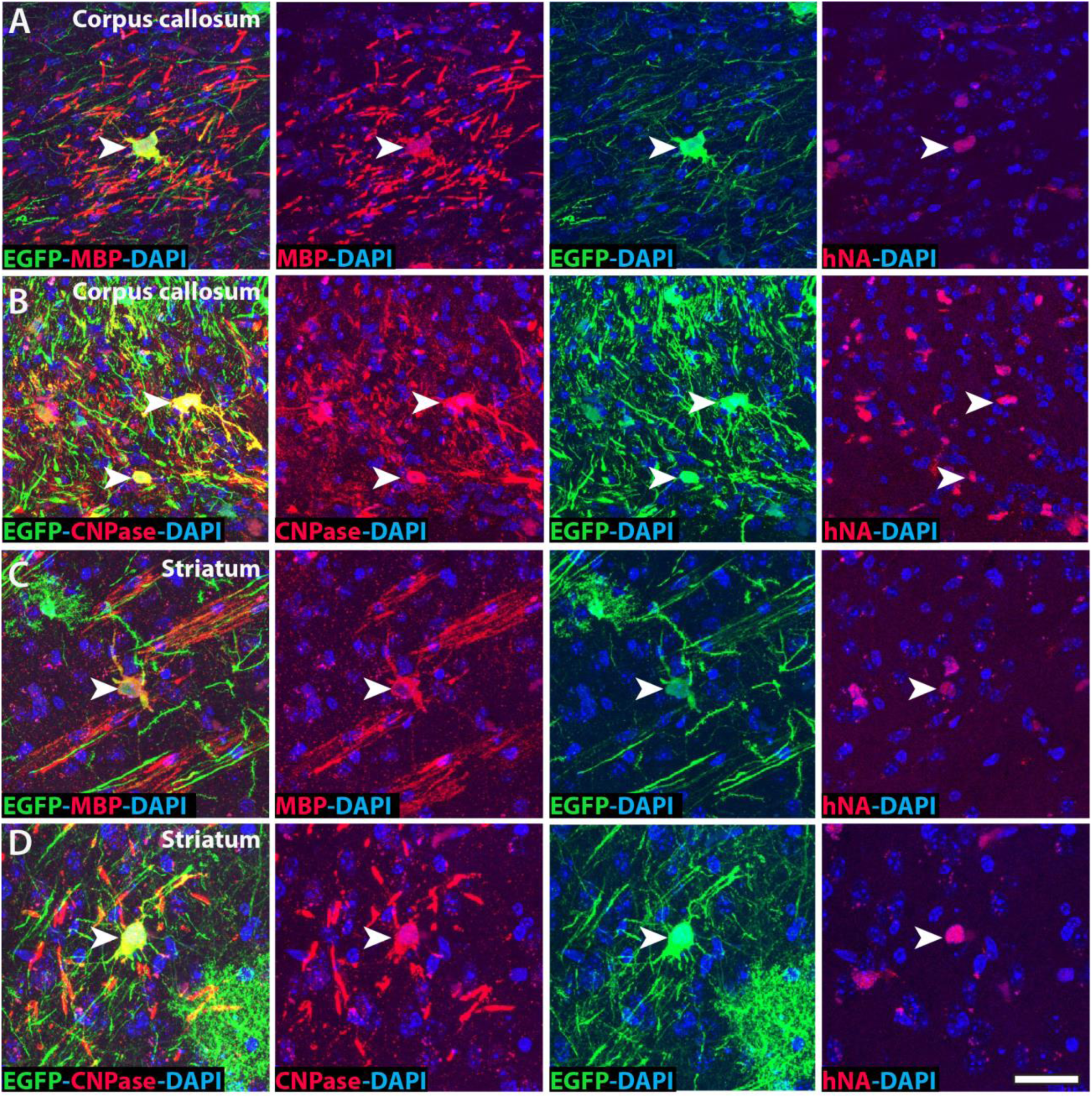
AAV-CM1 infects myelinated oligodendrocytes as well as their progenitors. AAV-CM1 exhibited efficient infection of mature myelinating oligodendrocytes, as well as of hGPCs. In this example, AAV5-CM1 was delivered intracisternally, after induction of systemic hypertonicity, in 16-week-old human glial chimeric shiverer mice, in which most forebrain myelinating oligodendrocytes and glial progenitors are of human origin. When sacrificed 3 weeks later, robust infection of CNPase (cyclic nucleotide diphosphatase)-defined and myelin basic protein (MBP)-defined myelinated oligodendrocytes – as well as parenchymal astrocytes and persistent hGPCs - was noted in the callosal (**A-B**) and corticostriatal (**C-D**) white matter tracts.

## DISCUSSION

Glial progenitor cells comprise an especially important phenotype of the adult human CNS, as these cells can give rise to both oligodendrocytes and astrocytes; under select circumstances they may also give rise to neurons as well. As such, this is an important phenotype to be able to target with gene therapeutics, both efficiently and specifically so. While AAVs have been evolved to target a number of cell types of the adult central nervous system, most such efforts have focused on mouse cells; among those studies focused on human cells, most were necessarily conducted in vitro, as few systems appropriate for targeting human brain cells in vivo have been developed. In this work, we report the development of a new set of AAVs, evolved from AAV serotype AAV5, that selectively and efficiently infect human glial progenitor cells (hGPCs) and their derivatives in vivo, with little off-target CNS or systemic infection. To do so, we selected candidate viruses in vivo, introducing libraries of randomly mutagenized, Cre recombinase-reported AAV5s intracisternally into mice that had been neonatally chimerized with Cre recombinase-expressing hGPCs. This approach combined the methodology of the CREATE strategy, as first described by Deverman, Gradinaru and colleagues^6^, with human glial chimeric brains^30,38^ as our platform for in vivo viral selection. Critically, we achieved robust glial infection in these mice using delivery via the perivascular glymphatic system^24,40^, so as to avoid the blood-brain barrier and hence enable efficient parenchymal entry of our engineered viruses, using a clinically feasible strategy.

The hGPCs in these chimerized mice were derived from hESCs that had been edited to express Cre recombinase under the regulatory control of the PDGFRA promoter, so that only human GPCs – which specifically express PDGFRα^27^ – expressed Cre. We injected the resultant human glial chimeras intracisternally with a library of AAVs, each of which expressed a random heptapeptide insertion into an AAV binding domain that regulates target cell adhesion. Each harbored a BGH poly-adenylation sequencefollowed by an inverted SV40 polyadenylation sequence, flanked by distinct Lox sequences (Lox66 and Lox71), which upon recombination in PDGFRA-Cre expressing cells, allowed the amplification of those viruses that had successfully transduced human GPCs. This approach identified those variants that selectively infected hGPCs and their derivatives in vivo; the exclusion of some of these, based on their infection of host liver, then allowed the selection of gliotropic viruses that exhibited minimal infection of non-neural or other systemic phenotypes.

Of note, the persistent neuronal infection by these glial-evolved capsid variants was an expected result of our approach, While our capsids modifications included the addition of random sequences and selection for those that bound glia, we did not delete or mutagenize the remaining VP1 sequence or modify its other exposed loops, which are together responsible for its neuronal avidity. As such, we added significant glial tropism to these vectors, without necessarily removing neuronal infectivity; our filtration against other tissues (liver, spleen, kidney) thus excluded variants cross-reactive with non-CNS tissue, but still allowed variants that retained neuronal infectivity. Since many of our potential disease targets of interest involve both neuronal and glial pathology, this is a desirable feature in most instances. In some disorders though, glial specificity might be preferred. In these cases, further work to selectively remove the neuronal sequences on VP1 will be needed, to create vectors whose range of infectivity is strictly restricted to glia. Until then, in such cases where glial-specific gene expression to the exclusion of neurons is a priority, one may add cell type-selective regulatory elements – enhancers and/or promoters - to further specify gene expression to glial phenotypes of interest. In this regard, a number of enhancers have been identified that can direct gene expression to astrocytes and/or oligodendrocytes^41^, whose pairing with capsids permitting cell type-selective infection may provide especially robust vectors for safe clinical use.

To provide these vectors to the brain in as widespread and efficient a manner possible, while minimizing dose and systemic spread, we developed a multifaceted approach to enhance their glymphatic delivery. First, we combined AAV administration with systemic hypertonic saline, a strategy previously shown to improve CNS delivery of antibodies by transiently increasing CSF influx along perivascular spaces ^25,42,43^. In addition, we positioned the mice in the Trendelenburg position post-injection, leveraging gravity to facilitate rostral distribution of the vector. This method aligns with findings by Castle and colleagues ^44^, who demonstrated that maintaining rodents in Trendelenburg for two hours after intrathecal AAV9 infusion significantly increased gene transfer to cortical and basal forebrain regions, with a >15-fold increase in neuronal transduction relative to upright controls. Our combination of intracisternal viral delivery, systemic hyperosmotic modulation, and strategic physical positioning significantly and substantially enhanced infection throughout the brain parenchyma. Of note, our approach complements earlier studies utilizing AQP4 knockout mice, which exhibit impaired glymphatic clearance, leading to the increased retention of AAV vectors within the brain following intracerebroventricular administration ^45^. Together, these findings underscore the importance of integrating physiological manipulations with delivery strategies, so as to optimize CNS-targeted gene therapy.

These vectors, especially as combined with this delivery approach, may find use in a broad set of clinical indications, as an increasing numbers of neurological disorders become recognized as either glial in etiology, or with attendant and contributory glial pathology. These include the adult myelin disorders, such as progressive multiple sclerosis and age-related white matter loss, for which hGPCs and their oligodendrocytic progeny are clear targets. In addition, our hGPC-targeted viruses may selectively infect neoplastic stem and progenitor cells within malignant glial tumors, since human glioma cells may arise from resident hGPCs, and express high levels of the PDGFA receptor ^46^. Most broadly though, these selectively gliotropic vectors may find value in treating the neurodegenerative disorders, given the importance of astrocytic and oligodendroglial pathology in diseases as diverse as Huntington’s, schizophrenia, Parkinson’s, and Alzheimer’s ^47–50^ (reviewed in ^51^). For each of these, potential transcriptional targets have been identified, but the means of selectively and safely transducing a high proportion of the relevant glia in any given brain region in vivo – much less throughout the entire brain and CNS - have not previously been available. Here, we describe a vector selection platform that enables the production of AAV5s targeting human phenotypes of interest in vivo, while validating a glymphatic delivery pathway that both minimizes extra-cerebral viral spread and efficiently circumvents the blood-brain barrier, potentially enabling the delivery of therapeutic transgenes to brain glia in a clinically safe and effective manner.

## METHODS

### Generation of PDGFRA-Cre hESCs

The human embryonic stem cells (hESC) line GENEA019 (GENEA Biocells, Sydney, Australia; a female line), was used to generate hGPCs in this study. First at the hESC stage, Cre recombinase was inserted immediately downstream of the PDGFRA coding sequence and an intra-ribosomal entry site (IRES), the latter to allow bicistronic expression. The knock-in was performed by HDR-mediated insertion of the donor DNA, with the sgRNA targeting the PDGFRA gene using the following sequence, 5’-ctgtaactggcggattcgag-3’. This was targeted immediately after the stop of the coding sequence (**Figure 1A**), and the sgRNA was inserted downstream of a U6 promoter in the pSpCas9(BB)-2A-Puro (PX459) plasmid (gift from Feng Zhang; Addgene 62988)^28^. This plasmid contains spCas9 expressed under a CBH promoter. Both the donor and the guide/Cas9 plasmid were electroporated into undifferentiated GENEA019 hESCs, as we have previously described ^52^. The cells were selected with puromycin (0.5µg/ml; Thermo Fisher, A1113803) until colonies were large enough to be picked up for expansion and genotyping. The genomic integrity of selected homozygous clones was assessed via karyotype and array chromosomal genome hybridization (aCGH), and a lack of major structural variants was confirmed before hGPC production, all as described ^29^.

### Mouse Models

For screening viruses in wild-type, non-chimeric mice, we purchased C57BL/6J mice from Jax Laboratories. For screening viruses in human glial chimeras, with an emphasis on targeting hGPCs and astrocytes, we used Rag1^−/−^ immunodeficient mice (Taconic) chimerized neonatally with hESC GPCs; the brains of these mice typically exhibited widespread colonization hGPCs and their derived astrocytes.

For screening potential oligodendrocyte-tropic viruses, we used Rag2-null immunodeficient and myelin-deficient mice chimerized neonatally with hESC hGPCs. These mice reliably produced a largely humanized white matter. For this purpose, homozygous MBP^shi/shi^ shiverer mice (Jackson Laboratory) were bred with homozygous Rag2^−/−^ immunodeficient mice on the C3h background (Taconic) to produce myelin-deficient and immunodeficient mice, as described ^53^.

All mice were housed in a temperature and humidity-controlled environment (64-73°F, 30-70% humidity) within a pathogen-free colony room on a 12:12-hour light cycle. They were given ad libitum access to Modified ProLab RMH 3000, 5P00 containing 0.025% trimethoprim/0.124% sulfamethoxazole (Mod LabDiet 5P00), and autoclaved acid water (pH 2.5-3.0).

### AAV production

#### AAV capsid library construction and amplification

To generate a viral library, we modified the M-CREATE strategy ^6,7^. The plasmid to receive the 7-mer insert library was created using the backbone of the pAAV-MCS Expression Vector (Agilent; cat. VPK-410). This backbone, including AAV2 ITRs, was fused to an expression cassette, in which a portion of the AAV2-P40-Rep2 was included for maintaining the expression of the capsid proteins and their proper splicing. The capsid sequence was modified by inserting a GCT codon (Alanine) between Q574 and S575 of the AAV5 VP1 capsid to introduce an NheI cut site for insertion of a random Heptamer, as well as a silent mutation at L543 to destroy one of the XhoI sites and make it unique. The library was then created by amplifying the region between NheI and XhoI with AAV5-Cap-XhoI-Forward primer (5’-ccacgaccaataggatggagc-3’) and AAV5-Cap-NNN-NheI-Reverse primer (5’-gggcagtggtggagctagcNNNNNNNNNNNNNNNNNNNNNactagcctggttgttggtggc-3’). The PCR amplicon was then inserted into the capsid opened NheI and XhoI. Downstream of the Capsid gene, the construct also included two polyadenylation sequences: BGH (sense) and SV40 (anti-sense), both flanked by Lox66 and Lox71, to allow for a Cre-dependent single flip. The resulting library preps contained 4×10^5 variants by limiting dilution of the bacterial clones.

#### AAV2-Rep construct

To abolish the expression in cis of the native capsid and generate viral particles exclusively from the engineered capsid library, pAAV2/5 (Addgene 104964, donated by Melina Fan) was subjected to PCR-directed mutagenesis followed by recombination-mediated ligation (InFusion, Takara). In this construct (pAAV2/5-Rep2-ΔCap5-3STOP), the expression of Cap proteins was abrogated by introduction of nonsense mutations at the start of each of VP1 (Glu12Ter), VP2 (Lys6Ter), and VP3 (Gln14Ter). All cloning was performed using Takara Bio’s In-Fusion Snap Assembly Kit (Cat #638952).

#### Generating capsids with evolved peptide inserts

After enriched capsid variants were recovered, their 21 bp sequences were ordered as primers and ligated into AAV5 Rep-Cap containing the NheI and XhoI cut sites to create pAAV2/5-574-Empty-Recipient plasmid. Individual 21 bp, heptamer coding sequences for each viral variant (i.e., CM1 or CM6) were then inserted by PCR at the NheI end and ligated to the vector open NheI/XhoI, to creates a separate rep-cap plasmid for each variant.

#### Viral Amplification

For amplification of the final AAV library preps, HEK293FT cells were cultured in a 5% CO2 incubator at 37°C using 10% fetal bovine serum (FBS) formulated in high glucose DMEM media (Thermo Fisher). Previously published protocols ^54^ were followed to produce the AAVs. Briefly, per viral prep, ten 150-mm plates of approximately 80% confluency HEK293FT cells were transfected with 0.4 mg of DNA-mix containing the Capsid, Helper (Addgene #112867, a gift from James Wilson), and AAV shuttle plasmids (pAAV-CAG-EGFP, pAAV-CAG-FLEX-EGFP and pAAV-CAG-FLEX-tdTomato, all gifts from Edward Boyden, as Addgene #37825, #28306 and #28304, respectively), at a ratio of 0.6:0.3:0.1, along with PEI transfection reagent at a ratio of 1:4. The media was replaced at 24 and 72 hrs post-transfection and the cells and media were harvested at 120 hrs. At both the 72 and 120 hrs timepoints, the media was treated with polyethylene glycol (PEG) for precipitating the virus. Cell pellets were lysed using Salt Active Nuclease, and the viral particles were separated by iodixanol density gradient solution spun using Type 60Ti rotor, at 59,000 rpm for 3h. The resulting viral band was further washed on an Amicon filter (MW cut-off 30 kDa; Millipore, UFC910024). The virus was then aliquoted and stored at −80°C. Viral titers were established by way of a qPCR AAV titer kit (ABM, G931), as per the manufacturer’s instructions.

### Production of hGPCs and establishment of human glial chimeric mice

Glial progenitor cells were produced from PDGFRA-Cre transduced human ESCs (GENEA-019; Genea Biocells) or from unmodified hESCs (WA09 line, WiCell Research Institute) using our previously described protocol ^29^. At 130-140 DIV, hGPCs were collected in Ca^2+^/Mg^2+^-free Hanks’ balanced salt solution (HBSS^−/−^; Thermo Fisher, 14170112), then mechanically dissociated to small clusters by gentle pipetting, spun, and resuspended in cold HBSS^−/−^ at 1 × 10^5^ cells/µl. Newborn mice (postnatal day 1) received bilateral intrastriatal injections of 1 µl cell suspension, i.e., 1×10^5^ hGPCs/hemisphere, under cryoanesthesia, as we have described^29,30,38^. The injection tracks permitted donor cell infiltration and chimerization of the overlying corpus callosum and cortex as well.

### Cisternal viral delivery with systemic hypertonicity

At 12 weeks of age, the neonatally-chimerized mice were anesthetized with ketamine (100 mg/kg) and xylazine (10 mg/kg) and monitored for anesthetic depth via cessation of toe pinch reflex. Mice were affixed in a stereotaxic frame and had their cisterna magna exposed. Surgical procedures were conducted as previously described ^26^. A 30-gauge cannula, connected by PE10 tubing to a 100 µl Hamilton syringe (gas-tight 1700 series), was implanted into the cisterna magna and secured with cyanoacrylate glue and dental cement. 10 µl of high titer (>10^12^ gc/ml) AAV injectate was then infused into the cisterna magna at 2 µl/min using a syringe micropump (Harvard Apparatus). Concurrent with infusion start, the mice were administered 20ul/g body weight IP injection of 1M hypertonic saline as previously described ^55^. Once the infusion was complete, mice were removed from the stereotaxic frame and placed in a prone Trendelenburg position on a −45-degree slope for 30 min, then returned to their home cage. Anesthetic depth was maintained for 1 hour following the time at which the infusion began. Animals were given analgesic and allowed to recover for either 7 or 21 days, depending on the experiment, at which time the mice were anesthetized and sacrificed via transcardiac perfusion with 20 ml of cold phosphate buffered saline (PBS) followed by 20 ml 4% paraformaldehyde (PFA). Tissues were then harvested, and immersion fixed in 4% PFA for an additional two hours.

### AAV capsid identification

One week post injection, mice were sacrificed and livers, corpora callosa, and striata dissected. Genomic DNA was then extracted from either liver or pooled callosal and striatal cells using the Wizard SV Genomic DNA Purification System (A2361). Next Generation Sequencing Libraries were generated by using a nested PCR and PrimeSTAR GXL DNA Polymerase (Takara Bio, R050A) to recover either recombined or non-recombined viral genomes from the samples. This PCR consisted of 10 cycles of outer PCR after which primers with added Illumina adapters and sample indices were spiked into the reactions and run for 25 more cycles to give each sample a unique barcode. These were run on a 1% agarose gel and the expected 352 bp band was purified. Final samples were indexed using NexteraXT Indexing and sequencing on an Illumina MiSeq V2 Nano (150bp paired end).

Following sequencing and demultiplexing all subsequent analysis was carried out in R; the 21bp DNA sequences, corresponding to the random heptamer coding sequences, were extracted based on their flanking sequences. Each sample was then normalized by sequencing depth and converted into 7aa peptide sequences. Within individual animal libraries, recombined brain libraries were then evaluated for enrichment vs non-recombined brain or liver libraries. From each mouse, the top 5-most enriched human-selective heptamers were used to generated viral particles for downstream testing. All code for this analysis may be found at: https://github.com/CTNGoldmanLab/hGPC_AAV_Capsid_Evolution.

### Tissue processing and immunohistochemistry

Mice were transcardially perfused with HBSS (Thermo Fisher Scientific) followed by 4% paraformaldehyde (PFA) in 0.1M PBS at 3 weeks following AAV delivery. The brains, livers, spleens and kidneys were dissected, post-fixed in 4% PFA for 2h, and stored in 1x PBS. Brains were cryopreserved in 0.1M PBS + 30% sucrose before embedding in OCT. Cryopreserved brains were sagittally sectioned into 20 μm-thick serial sections using a cryostat.

Brain sections were blocked with 5% normal goat serum in 0.1 M PBS + 0.3% Triton-X for 1 hr at RT and incubated overnight at 4°C with primary antibodies against EGFP (chicken, 1:1000, Thermo Fisher Scientific), human nuclear antigen (mouse, 1:400, Abcam ab254080), PDGFRα (rabbit, 1:600, Cell Signaling Technology), Olig2 (rabbit, 1:500, Millipore), NeuN (rabbit, 1:800, Millipore), Sox9 (rabbit, 1:500, Abcam), MBP (1:400, Abcam ab40390), anti-dtTomato (1:400, Invitrogen, M11217), CNPase (1:800, Abcam ab18527), NeuN (1:400, Millipore ABN78MI), or HA tag (mouse, 1:400, Invitrogen). Immunostaining was visualized with species-specific secondary antibodies conjugated with AlexaFluor-488, AlexaFluor-568 (1:400, Invitrogen) or AlexaFluor-647 (1:200, Invitrogen). Lastly, sections were coverslipped with mounting media containing DAPI (Vector Laboratories). Slides were sealed and imaged with Olympus Slideview vs200 or a Leica DM6000B confocal microscope

### Quantitative PCR

Sample tissue from spleen, kidney or liver were homogenized and DNA extracted using the Wizard® Genomic DNA Purification Kit following the manufacturer protocol (Promega). qPCR was run using TaqMan probes for the 18s subunit ribosomal gene and EGFP (ThermoFisher, IDs Mr04329676 and Mm03928990 respectively). Data were analyzed in R as one-way ANOVAs with Tukey post hoc tests and visualized using ggplot2.

### Fluorescence in situ hybridization (FISH)

One-year-old mice engrafted at PND1 with G19-Cre GPCs were anesthetized and rapidly decapitated. Brains were then harvested, frozen in chilled isopentane, embedded in OCT (TissueTek), sectioned sagittally at 10 μm on a cryostat and stored at −80°C. Slides with tissue sections were immersed in 4% PFA for 15 min, rinsed twice with 1X PBS, and dehydrated at room temperature in increasing ethanol solutions: 50%, 70%, 100%, 100% for 5mins each. Using the RNAscope® Multiplex Fluorescent Reagent Kit (323100, ACDBio), tissue sections were incubated in hydrogen peroxide for 10 min in a humidified box, rinsed with distilled water, and incubated with protease III for 30mins at room temperature. Hybridization probes against Cre recombinase (312281-C1, ACDBio), human PDGFRa (604481-C2, ACDBio), and human Olig2 (424191-C3, ACDBio) or human NeuN (415591-C3, ACDBio) were then incubated for 2hrs at 40°C. Control probes were also incubated in separate tissue sections. Probe signals were then serially amplified for 30mins at 40°C each and individually developed starting with Opal 520 (1:750; FP1487001KT, PerkinElmer), followed by Opal 570 (1:750; FP1488001KT, PerkinElmer), then Opal 690 (1:1500; FP1497001KT, PerkinElmer). Slides were coverslipped with Vectashield with DAPI for imaging.

## ACKNOWLEDGEMENTS

This work was supported by CNS2, Inc., the Adelson Medical Research Foundation, and NIH grants R01AG072298 and R01NS110776. We thank Christina Trojel-Hansen for advice and critical comments. We are grateful to Victoria Bermeo Vargas and Thulasi Ram for histological support, and to Paul Tobin and Ariba Khan for assistance in quantification. We thank the University of Rochester Genomics Research Center for its assistance in sequencing and analysis.

## AUTHOR CONTRIBUTIONS

AC and AB designed and constructed the viral vectors, and NW and WB raised and purified them; EN and AC conducted the intracisternal injections and optimized the glymphatic delivery approach together with MN; Ab and DK engineered the PDGFRA-Cre hESCs, and XL and DCM produced hGPCs from them; SS established the chimeric mice; WB performed the QPCR; AC, EN, JD, and AI performed the immunohistology and quantitative imaging, and JC the in situ hybridizations for CRE; RS and JNM analyzed the PCR and genomic data; AB, MN and SAG designed the study and analyzed all data; SAG wrote the paper, with input from all co-authors.

## EXTENDED DATA

**Extended Data Figure 1 (related to Figure 1).**
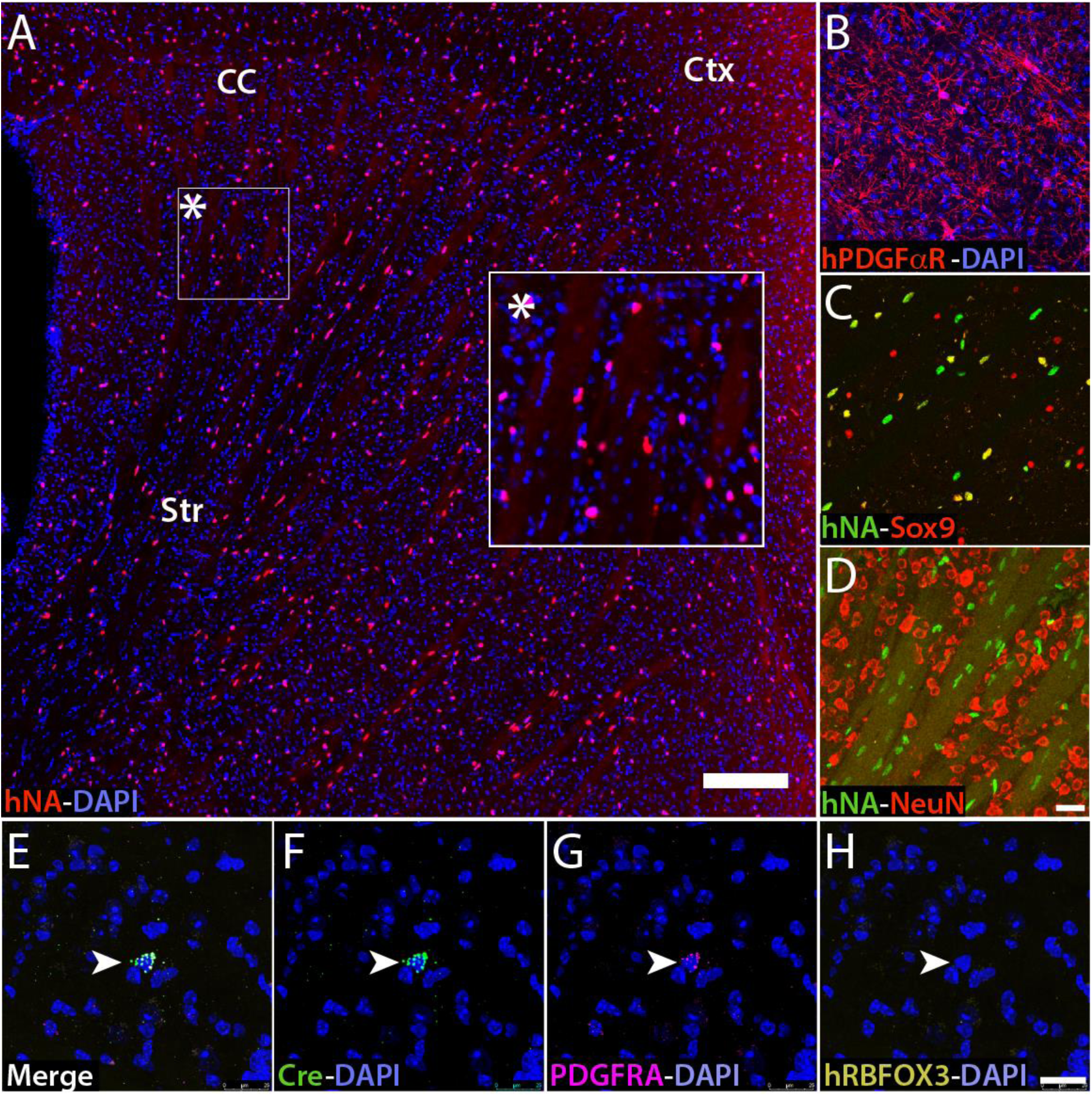
Colonization of mouse forebrain with human PDGFRA:Cre glia allows in vivo capsid selection. **A.** Mice neonatally engrafted with hESC-derived PDGFRA:Cre-expressing glial progenitor cells (hGPCs) displayed robust colonization of the striatum by 14 weeks post-transplant. Human donor-derived cells either remained as hGPCs (**B**), or differentiated into Sox9-expressing astrocytes (**C**), but did not differentiate into neurons (**D**). **E.** RNAscope multiplex in situ hybridization on striata of engrafted mice showed that the CRE-specific probe (**F**) colocalized with PDGFRA (**G**) but not with that of NeuN (**H**). Scale: 100 µm (**A**); 25 µm (**D**, **E-H**).

**Extended Data Figure 2 (related to Figure 1).**
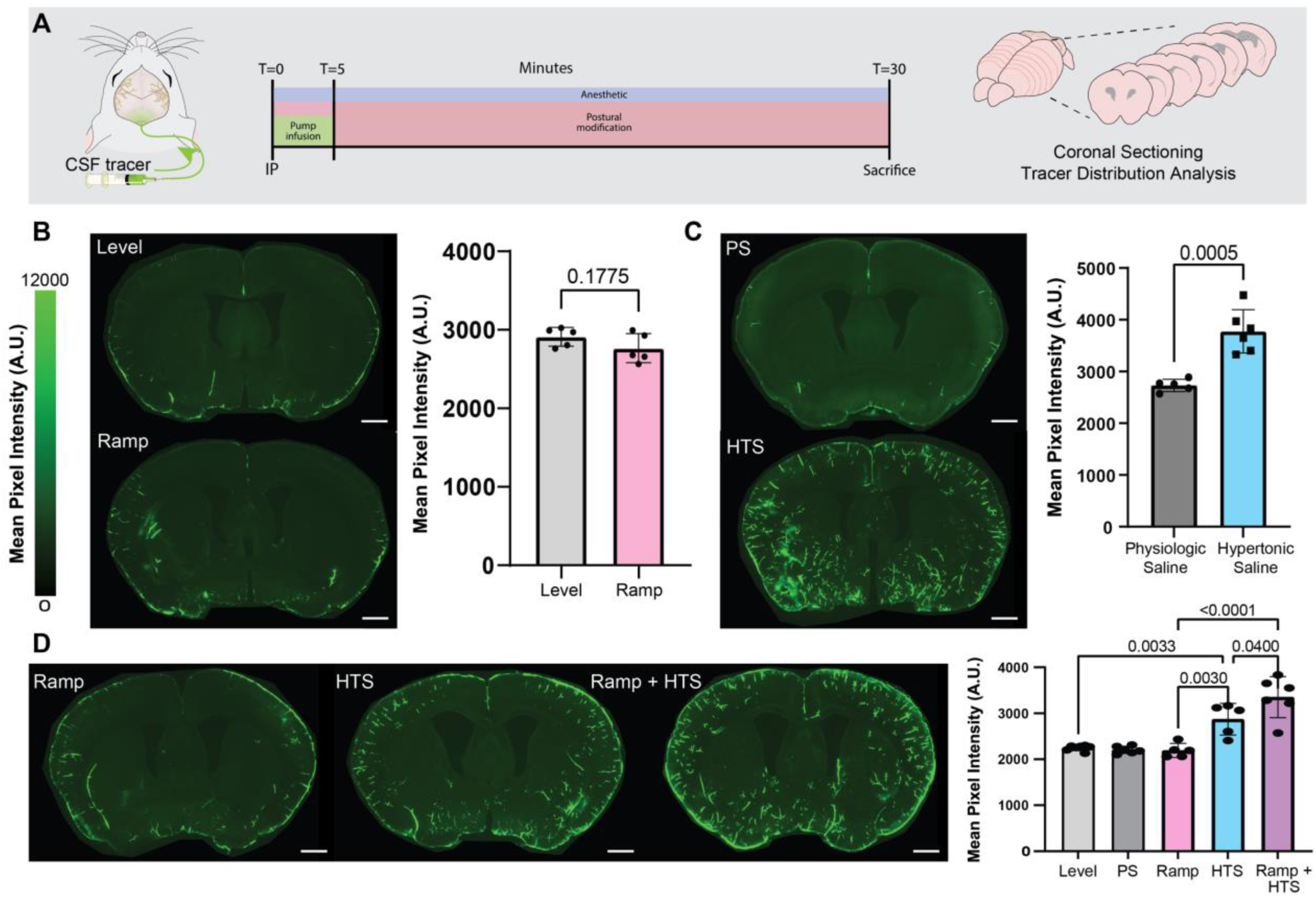
Hypertonic intracisternal delivery of tracer achieves widespread brain access. **A.** Experimental schema of cisterna magna CSF tracer infusions (FITC-dextran, 2,000kDa, 2.5% w/v in saline; 10 µl delivered at 2 µl/min), done under ketamine/xylazine anesthesia with postural modification, osmotic manipulation via hypertonic saline (1M, 20 µl/g), or both. **B.** Representative images of coronal brain slices with CSF tracer influx (2000kDa FITC-dextran) after being placed prone on a level surface (left, top) or a declined plane (left, bottom) until t = 30 minutes post infusion start. (Quantification of fluorescent intensity of coronal sections shows no change in influx with postural modification alone (right, p=.1775). **C.** Representative coronal brain slices with CSF tracer influx (2000kDa FITC-dextran) receiving either physiologic saline (PS, *top*) or hypertonic saline (HTS, *bottom*). Quantification of fluorescent intensity shows increased CSF tracer influx with hypertonic saline treatment (*right panel,* p=.0005; PS, n=5 mice, HTS, n=6). **D.** Representative images of coronal brain slices with 2000kDa FITC-dextran influx comparing injection in prone Trendelenburg position on a declined plane (*left*), vs. hypertonic saline alone (*middle*), or both (*right*). Quantification of fluorescent intensity shows increased influx with HTS and further increase when HTS is combined with postural modification. **B-C**, unpaired Students t-tests. **D**, One-way ANOVA, post hoc p values as noted. For **B-D**, individual mean and standard deviation were plotted, and each data point is an individual mouse (n= 5 level, 5 ramp, 5 PS, 6 HTS, 6 Ramp + HTS). Ramp = postural modification, HTS = hypertonic saline, PS = physiologic saline. Scale: 1 mm.

**Extended Data Figure 3 (related to Figure 1).**
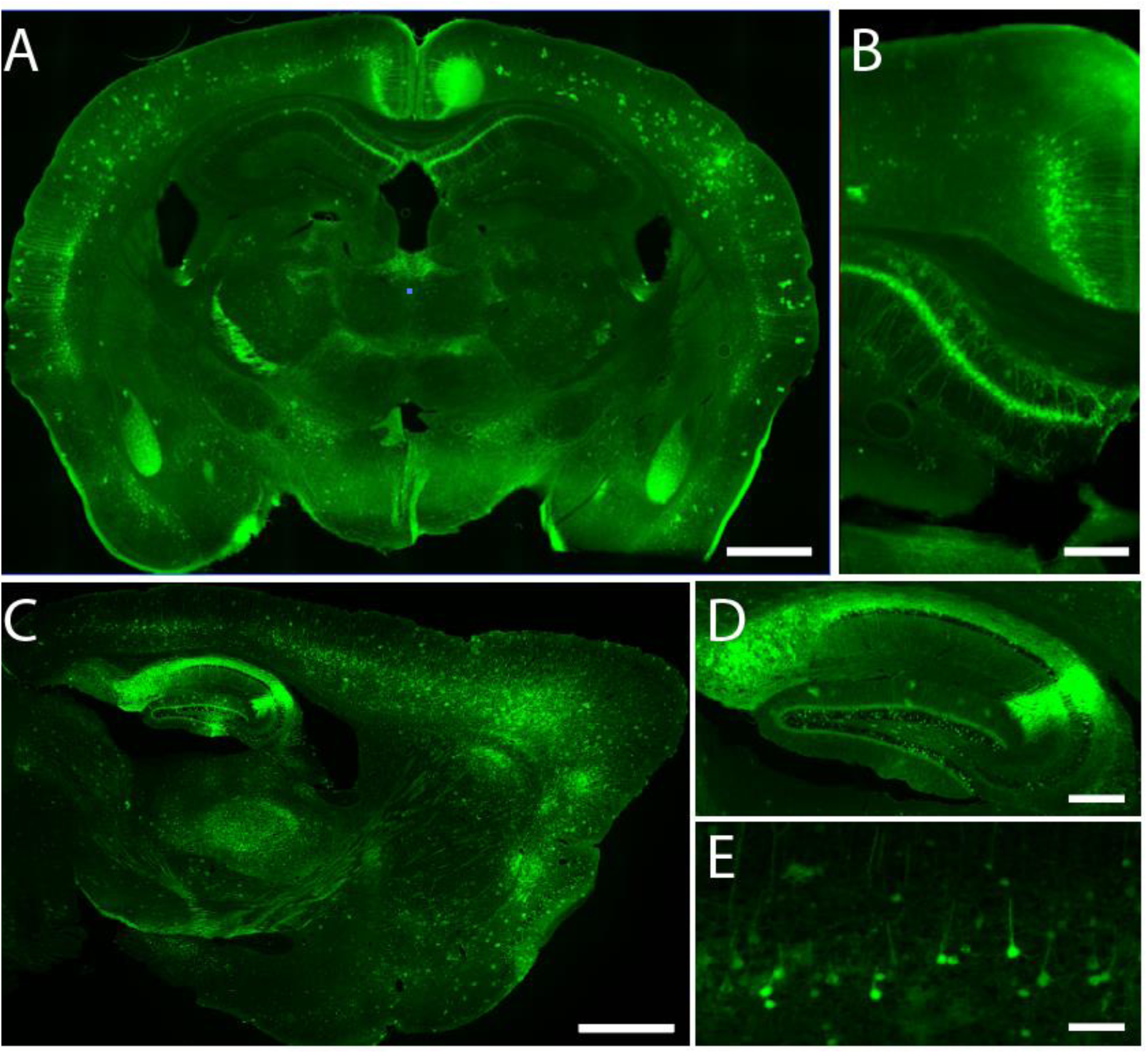
Intracisternally-delivered AAV pervades the forebrain parenchyma via glymphatic spread. To establish the efficacy of our intracisternal hypertonic delivery protocol using a well-described control vector before testing our modified capsid variants, we injected a set of adult mice with either AAV5-retro (**A-B**) or WT AAV5 (**C-E**). For the AAV5-retro injections, of **A-B**, we inserted the neurotropic retro cassette (encoding the peptide LADQDYTKTA) in between Q574 and S575 of the AAV5 VP1 capsid protein; this locus presents an exposed sialic acid-binding loop domain by which target specificity may be modified. A single intracisternal injection under systemic hypertonicity yielded high-efficiency infection of forebrain hippocampal pyramidal and selected corticofugal neurons, as has been described for parenchymally-administered AAV2-based AAV retro^34^. **C-E**, Similarly, intracisternal WT AAV5 control virus yielded predominantly neuronal labeling thought the forebrain, most prominently so in the hippocampus (**D**) and cortex (**E**). These results confirm that glymphatic-mediated intracisternal delivery supports efficient, brain-wide parenchymal access and neuronal gene transfer. Scale: **A, C**: 1 mm; **B, D**: 250 µm; **E**: 100 µm.

**Extended Data Figure 4 (related to Figure 2).**
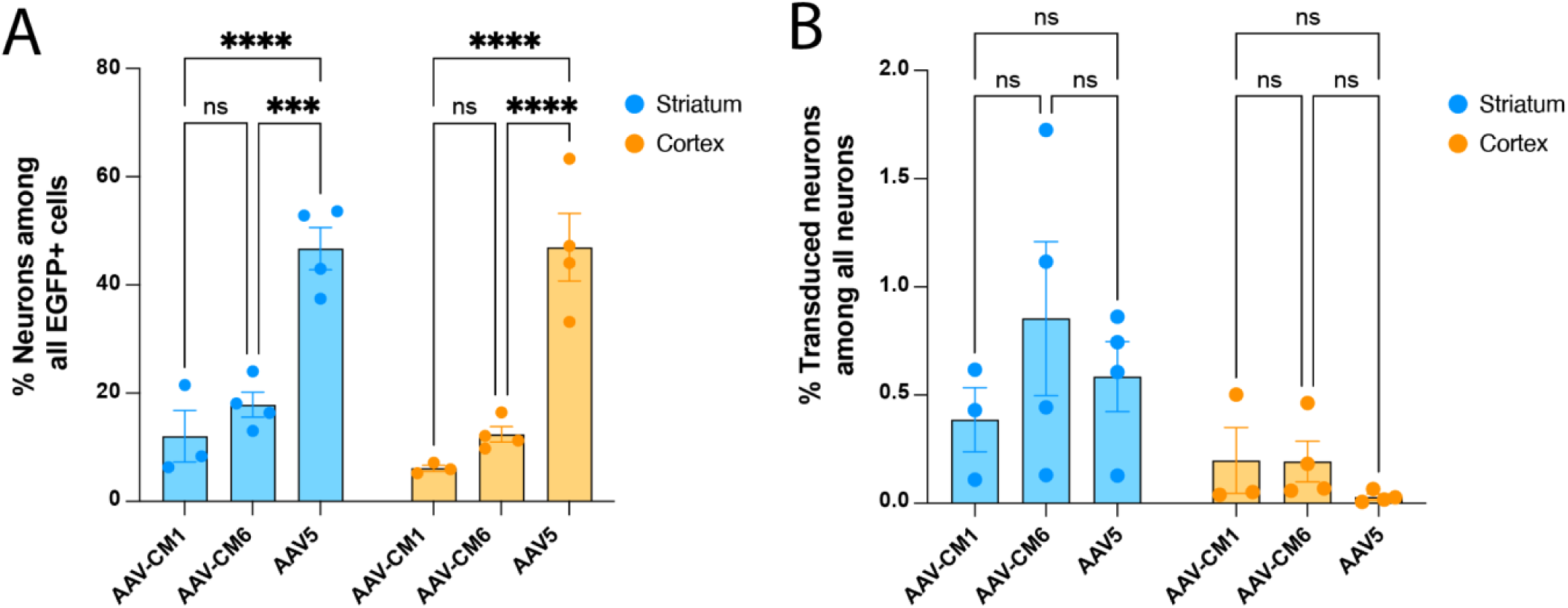
Adventitious neuronal infection persists in gliotropic variants. Despite their strong relative gliotropism, some neuronal transduction persisted to a variable extent in each of these engineered viral variants. **A,** Both AAV-CM1 and AAV-CM6 exhibited significantly less neuronal infection than the unmodified AAV5s from which they were derived, in both the neostriatum and the overlying neocortex. In contrast, in controls injected with unmodified AAV5 almost 50% of infected cells were neurons. **B,** In the same brains and regions, less than 1% of all neurons were infected, and as noted in **A**, these comprised <20% of all cells infected by either CM1 or CM6. 2-way ANOVA with Tukey’s post-hoc tests. Mean ± SEM. ns p > 0.05, ***p < 0.001, ****p < 0.0001.

**Extended Data Figure 5 (related to Figure 2).**
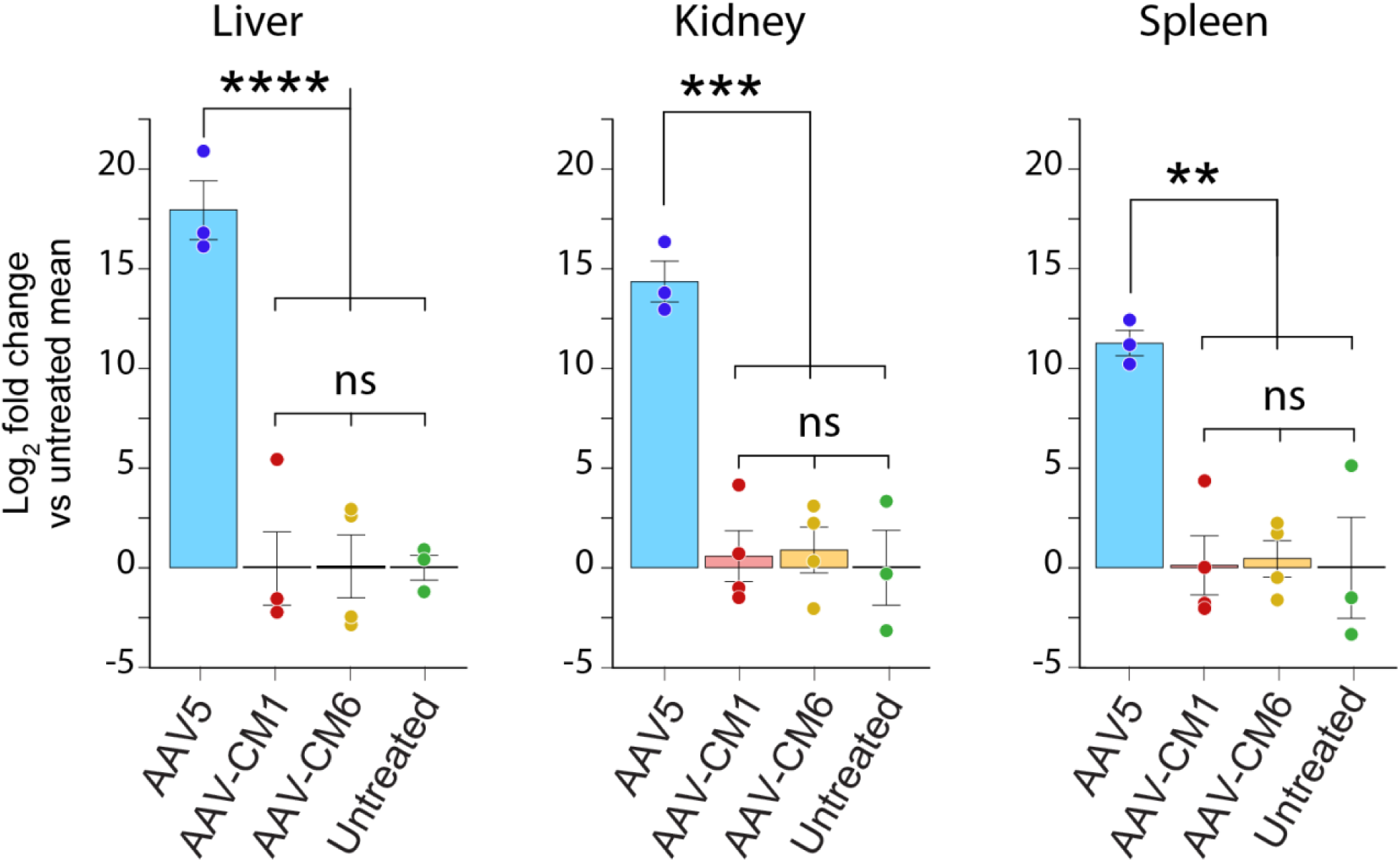
Selected capsid-evolved AAVs manifest little systemic infection. Using PCR to assess the hepatic viral genome concentrations (vg/g) allowed estimation of the degree of off-target infection by each capsid variant. Only those candidates lacking appreciable hepatic infection were developed further. **A-C.** When assessed a week after intracisternal delivery, two capsid candidates that survived this initial filtration, AAV-CM1 and AAV-CM6, exhibited little or no infection of the liver **(A**), or of the kidney (**B**) or spleen (**C**), suggesting the strong phenotypic selectivity afforded by capsid evolution on hGPCs in vivo. One-way ANOVA with Tukey post hoc tests. Means ± SEM. **p<0.01, ***p<0.001; ****p<0.0001.

## REFERENCES

1. Franklin, R.J.M., Bodini, B., and Goldman, S.A. (2024). Remyelination in the Central Nervous System. Cold Spring Harbor perspectives in biology 16. 10.1101/cshperspect.a041371.

2. Groh, J., and Simons, M. (2025). White matter aging and its impact on brain function. Neuron 113, 127–139. 10.1016/j.neuron.2024.10.019.

3. Huynh, N.P.T., Osipovitch, M., Foti, R., Bates, J., Mansky, B., Cano, J.C., Benraiss, A., Zhao, C., Lu, Q.R., and Goldman, S.A. (2024). Shared patterns of glial transcriptional dysregulation link Huntington’s disease and schizophrenia. Brain. 10.1093/brain/awae166.

4. Osipovitch, M., Asenjo Martinez, A., Mariani, J.N., Cornwell, A., Dhaliwal, S., Zou, L., Chandler-Militello, D., Wang, S., Li, X., Benraiss, S.J., et al. (2019). Human ESC-Derived Chimeric Mouse Models of Huntington’s Disease Reveal Cell-Intrinsic Defects in Glial Progenitor Cell Differentiation. Cell Stem Cell 24, 107–122 e107. 10.1016/j.stem.2018.11.010.

5. Windrem, M.S., Osipovitch, M., Liu, Z., Bates, J., Chandler-Militello, D., Zou, L., Munir, J., Schanz, S., McCoy, K., Miller, R.H., et al. (2017). Human iPSC Glial Mouse Chimeras Reveal Glial Contributions to Schizophrenia. Cell Stem Cell 21, 195–208 e196. 10.1016/j.stem.2017.06.012.

6. Deverman, B.E., Pravdo, P.L., Simpson, B.P., Kumar, S.R., Chan, K.Y., Banerjee, A., Wu, W.L., Yang, B., Huber, N., Pasca, S.P., and Gradinaru, V. (2016). Cre-dependent selection yields AAV variants for widespread gene transfer to the adult brain. Nat Biotechnol 34, 204–209. 10.1038/nbt.3440.

7. Kumar, S.R., Miles, T.F., Chen, X., Brown, D., Dobreva, T., Huang, Q., Ding, X., Luo, Y., Einarsson, P.H., Greenbaum, A., et al. (2020). Multiplexed Cre-dependent selection yields systemic AAVs for targeting distinct brain cell types. Nat Methods 17, 541–550. 10.1038/s41592-020-0799-7.

8. Mandel, R.J., Marmion, D.J., Kirik, D., Chu, Y., Heindel, C., McCown, T., Gray, S.J., and Kordower, J.H. (2017). Novel oligodendroglial alpha synuclein viral vector models of multiple system atrophy: studies in rodents and nonhuman primates. Acta Neuropathol Commun 5, 47. 10.1186/s40478-017-0451-7.

9. Powell, S.K., Khan, N., Parker, C.L., Samulski, R.J., Matsushima, G., Gray, S.J., and McCown, T.J. (2016). Characterization of a novel adeno-associated viral vector with preferential oligodendrocyte tropism. Gene Ther 23, 807–814. 10.1038/gt.2016.62.

10. Nunes, M.C., Roy, N.S., Keyoung, H.M., Goodman, R.R., McKhann, G., Jiang, L., Kang, J., Nedergaard, M., and Goldman, S.A. (2003). Identification and isolation of multipotential neural progenitor cells from the subcortical white matter of the adult human brain. Nature medicine 9, 439–447.

11. Roy, N.S., Wang, S., Harrison-Restelli, C., Benraiss, A., Fraser, R.A., Gravel, M., Braun, P.E., and Goldman, S.A. (1999). Identification, isolation, and promoter-defined separation of mitotic oligodendrocyte progenitor cells from the adult human subcortical white matter. J Neurosci 19, 9986–9995. 10.1523/JNEUROSCI.19-22-09986.1999.

12. Scolding, N., Franklin, R., Stevens, S., Heldin, C.H., Compston, A., and Newcombe, J. (1998). Oligodendrocyte progenitors are present in the normal adult human CNS and in the lesions of multiple sclerosis. Brain : a journal of neurology 121 (Pt 12), 2221–2228. 10.1093/brain/121.12.2221.

13. Scolding, N.J., Rayner, P.J., and Compston, D.A. (1999). Identification of A2B5-positive putative oligodendrocyte progenitor cells and A2B5-positive astrocytes in adult human white matter. Neuroscience 89, 1–4.

14. Windrem, M.S., Nunes, M.C., Rashbaum, W.K., Schwartz, T.H., Goodman, R.A., McKhann, G., Roy, N.S., and Goldman, S.A. (2004). Fetal and adult human oligodendrocyte progenitor cell isolates myelinate the congenitally dysmyelinated brain. Nature medicine 10, 93–97.

15. Huang, Q., Chen, A.T., Chan, K.Y., Sorensen, H., Barry, A.J., Azari, B., Zheng, Q., Beddow, T., Zhao, B., Tobey, I.G., et al. (2023). Targeting AAV vectors to the central nervous system by engineering capsid-receptor interactions that enable crossing of the blood-brain barrier. PLoS biology 21, e3002112. 10.1371/journal.pbio.3002112.

16. Huang, Q., Chan, K.Y., Wu, J., Botticello-Romero, N.R., Zheng, Q., Lou, S., Keyes, C., Svanbergsson, A., Johnston, J., Mills, A., et al. (2024). An AAV capsid reprogrammed to bind human transferrin receptor mediates brain-wide gene delivery. Science 384, 1220–1227. 10.1126/science.adm8386.

17. Huang, Q., Chan, K.Y., Tobey, I.G., Chan, Y.A., Poterba, T., Boutros, C.L., Balazs, A.B., Daneman, R., Bloom, J.M., Seed, C., and Deverman, B.E. (2019). Delivering genes across the blood-brain barrier: LY6A, a novel cellular receptor for AAV-PHP.B capsids. PLoS One 14, e0225206. 10.1371/journal.pone.0225206.

18. Hinderer, C., Katz, N., Buza, E.L., Dyer, C., Goode, T., Bell, P., Richman, L.K., and Wilson, J.M. (2018). Severe Toxicity in Nonhuman Primates and Piglets Following High-Dose Intravenous Administration of an Adeno-Associated Virus Vector Expressing Human SMN. Hum Gene Ther 29, 285–298. 10.1089/hum.2018.015.

19. Arjomandnejad, M., Dasgupta, I., Flotte, T.R., and Keeler, A.M. (2023). Immunogenicity of Recombinant Adeno-Associated Virus (AAV) Vectors for Gene Transfer. BioDrugs 37, 311–329. 10.1007/s40259-023-00585-7.

20. Duan, D. (2023). Lethal immunotoxicity in high-dose systemic AAV therapy. Mol Ther 31, 3123–3126. 10.1016/j.ymthe.2023.10.015.

21. Gardin, A., and Ronzitti, G. (2023). Current limitations of gene therapy for rare pediatric diseases: Lessons learned from clinical experience with AAV vectors. Arch Pediatr 30, 8S46–48S52. 10.1016/S0929-693X(23)00227-0.

22. Guo, Y., Chen, J., Ji, W., Xu, L., Xie, Y., He, S., Lai, C., Hou, K., Li, Z., Chen, G., and Wu, Z. (2023). High-titer AAV disrupts cerebrovascular integrity and induces lymphocyte infiltration in adult mouse brain. Mol Ther Methods Clin Dev 31, 101102. 10.1016/j.omtm.2023.08.021.

23. Ling, Q., Herstine, J.A., Bradbury, A., and Gray, S.J. (2023). AAV-based in vivo gene therapy for neurological disorders. Nat Rev Drug Discov 22, 789–806. 10.1038/s41573-023-00766-7.

24. Iliff, J.J., Wang, M., Liao, Y., Plogg, B.A., Peng, W., Gundersen, G.A., Benveniste, H., Vates, G.E., Deane, R., Goldman, S.A., et al. (2012). A paravascular pathway facilitates CSF flow through the brain parenchyma and the clearance of interstitial solutes, including amyloid beta. Science translational medicine 4, 147ra111. 10.1126/scitranslmed.3003748.

25. Plog, B.A., Mestre, H., Olveda, G.E., Sweeney, A.M., Kenney, H.M., Cove, A., Dholakia, K.Y., Tithof, J., Nevins, T.D., Lundgaard, I., et al. (2018). Transcranial optical imaging reveals a pathway for optimizing the delivery of immunotherapeutics to the brain. JCI Insight 3. 10.1172/jci.insight.126138.

26. Xavier, A.L.R., Hauglund, N.L., von Holstein-Rathlou, S., Li, Q., Sanggaard, S., Lou, N., Lundgaard, I., and Nedergaard, M. (2018). Cannula Implantation into the Cisterna Magna of Rodents. J Vis Exp. 10.3791/57378.

27. Sim, F.J., McClain, C.R., Schanz, S.J., Protack, T.L., Windrem, M.S., and Goldman, S.A. (2011). CD140a identifies a population of highly myelinogenic, migration-competent and efficiently engrafting human oligodendrocyte progenitor cells. Nat Biotechnol 29, 934–941. 10.1038/nbt.1972.

28. Ran, F.A., Hsu, P.D., Wright, J., Agarwala, V., Scott, D.A., and Zhang, F. (2013). Genome engineering using the CRISPR-Cas9 system. Nat Protoc 8, 2281–2308. 10.1038/nprot.2013.143.

29. Wang, S., Bates, J., Li, X., Schanz, S., Chandler-Militello, D., Levine, C., Maherali, N., Studer, L., Hochedlinger, K., Windrem, M., and Goldman, S.A. (2013). Human iPSC-derived oligodendrocyte progenitor cells can myelinate and rescue a mouse model of congenital hypomyelination. Cell Stem Cell 12, 252–264. 10.1016/j.stem.2012.12.002.

30. Windrem, M.S., Schanz, S.J., Guo, M., Tian, G.F., Washco, V., Stanwood, N., Rasband, M., Roy, N.S., Nedergaard, M., Havton, L.A., et al. (2008). Neonatal chimerization with human glial progenitor cells can both remyelinate and rescue the otherwise lethally hypomyelinated shiverer mouse. Cell Stem Cell 2, 553–565. 10.1016/j.stem.2008.03.020.

31. Windrem, M.S., Schanz, S.J., Morrow, C., Munir, J., Chandler-Militello, D., Wang, S., and Goldman, S.A. (2014). A competitive advantage by neonatally engrafted human glial progenitors yields mice whose brains are chimeric for human glia. J Neurosci 34, 16153–16161. 10.1523/JNEUROSCI.1510-14.2014.

32. Iliff, J.J., Goldman, S.A., and Nedergaard, M. (2015). Implications of the discovery of brain lymphatic pathways. Lancet Neurol 14, 977–979. 10.1016/S1474-4422(15)00221-5.

33. Louboutin, J.P., Wang, L., and Wilson, J.M. (2005). Gene transfer into skeletal muscle using novel AAV serotypes. J Gene Med 7, 442–451. 10.1002/jgm.686.

34. Tervo, D.G., Hwang, B.Y., Viswanathan, S., Gaj, T., Lavzin, M., Ritola, K.D., Lindo, S., Michael, S., Kuleshova, E., Ojala, D., et al. (2016). A Designer AAV Variant Permits Efficient Retrograde Access to Projection Neurons. Neuron 92, 372–382. 10.1016/j.neuron.2016.09.021.

35. Castle, M.J., Turunen, H.T., Vandenberghe, L.H., and Wolfe, J.H. (2016). Controlling AAV Tropism in the Nervous System with Natural and Engineered Capsids. Methods Mol Biol 1382, 133–149. 10.1007/978-1-4939-3271-9_10.

36. Wang, Y., Yang, C., Hu, H., Chen, C., Yan, M., Ling, F., Wang, K.C., Wang, X., Deng, Z., Zhou, X., et al. (2022). Directed evolution of adeno-associated virus 5 capsid enables specific liver tropism. Mol Ther Nucleic Acids 28, 293–306. 10.1016/j.omtn.2022.03.017.

37. Schnutgen, F., Doerflinger, N., Calleja, C., Wendling, O., Chambon, P., and Ghyselinck, N.B. (2003). A directional strategy for monitoring Cre-mediated recombination at the cellular level in the mouse. Nat Biotechnol 21, 562–565. 10.1038/nbt811.

38. Windrem, M.S., Schanz, S.J., Morrow, C., Munir, J., Chandler-Militello, D., Wang, S., and Goldman, S.A. (2014). A Competitive Advantage by Neonatally Engrafted Human Glial Progenitors Yields Mice Whose Brains Are Chimeric for Human Glia. J. Neurosci. 34, 16153–16161. 10.1523/JNEUROSCI.1510-14.2014.

39. Windrem, M.S., Schanz, S.J., Zou, L., Chandler-Militello, D., Kuypers, N.J., Nedergaard, M., Lu, Y., Mariani, J.N., and Goldman, S.A. (2020). Human Glial Progenitor Cells Effectively Remyelinate the Demyelinated Adult Brain. Cell Rep 31, 107658. 10.1016/j.celrep.2020.107658.

40. Nedergaard, M., and Goldman, S.A. (2020). Glymphatic failure as a final common pathway to dementia. Science (New York, N.Y.) 370, 50–56. 10.1126/science.abb8739.

41. Mich, J.K., Sunil, S., Johansen, N., Martinez, R.A., Leytze, M., Gore, B.B., Mahoney, J.T., Ben-Simon, Y., Bishaw, Y., Brouner, K., et al. (2023). Enhancer-AAVs allow genetic access to oligodendrocytes and diverse populations of astrocytes across species. bioRxiv. 10.1101/2023.09.20.558718.

42. Salegio, E.A., Hancock, K., and Korszen, S. (2023). Pre-clinical delivery of gene therapy products to the cerebrospinal fluid: challenges and considerations for clinical translation. Frontiers in molecular neuroscience Volume 16 - 2023. 10.3389/fnmol.2023.1248271.

43. Chen, X., Lim, D.A., Lawlor, M.W., Dimmock, D., Vite, C.H., Lester, T., Tavakkoli, F., Sadhu, C., Prasad, S., and Gray, S.J. (2023). Biodistribution of Adeno-Associated Virus Gene Therapy Following Cerebrospinal Fluid-Directed Administration. Human gene therapy 34, 94–111. 10.1089/hum.2022.163.

44. Castle, M.J., Cheng, Y., Asokan, A., and Tuszynski, M.H. (2018). Physical positioning markedly enhances brain transduction after intrathecal AAV9 infusion. Sci Adv 4, eaau9859. 10.1126/sciadv.aau9859.

45. Murlidharan, G., Crowther, A., Reardon, R.A., Song, J., and Asokan, A. (2016). Glymphatic fluid transport controls paravascular clearance of AAV vectors from the brain. JCI insight 1, e88034. 10.1172/jci.insight.88034.

46. Persson, A.I., Petritsch, C., Swartling, F.J., Itsara, M., Sim, F.J., Auvergne, R., Goldenberg, D.D., Vandenberg, S.R., Nguyen, K.N., Yakovenko, S., et al. (2010). Non-stem cell origin for oligodendroglioma. Cancer Cell 18, 669–682. 10.1016/j.ccr.2010.10.033.

47. Depp, C., Sun, T., Sasmita, A.O., Spieth, L., Berghoff, S.A., Nazarenko, T., Overhoff, K., Steixner-Kumar, A.A., Subramanian, S., Arinrad, S., et al. (2023). Myelin dysfunction drives amyloid-β deposition in models of Alzheimer’s disease. Nature 618, 349–357. 10.1038/s41586-023-06120-6.

48. Huynh, N.P.T., Osipovitch, M., Foti, R., Bates, J., Mansky, B., Cano, J.C., Benraiss, A., Zhao, C., Lu, Q.R., and Goldman, S.A. (2024). Shared patterns of glial transcriptional dysregulation link Huntington’s disease and schizophrenia. Brain 147, 3099–3112. 10.1093/brain/awae166.

49. Kenigsbuch, M., Bost, P., Halevi, S., Chang, Y., Chen, S., Ma, Q., Hajbi, R., Schwikowski, B., Bodenmiller, B., Fu, H., et al. (2022). A shared disease-associated oligodendrocyte signature among multiple CNS pathologies. Nature Neuroscience 25, 876–886. 10.1038/s41593-022-01104-7.

50. Nasrabady, S.E., Rizvi, B., Goldman, J.E., and Brickman, A.M. (2018). White matter changes in Alzheimer’s disease: a focus on myelin and oligodendrocytes. Acta Neuropathol Commun 6, 22. 10.1186/s40478-018-0515-3.

51. Goldman, S.A. (2020). Glial evolution as a determinant of human behavior and its disorders. Ann N Y Acad Sci. 10.1111/nyas.14372.

52. Vieira, R., Mariani, J.N., Huynh, N.P.T., Stephensen, H.J.T., Solly, R., Tate, A., Schanz, S., Cotrupi, N., Mousaei, M., Sporring, J., et al. (2024). Young glial progenitor cells competitively replace aged and diseased human glia in the adult chimeric mouse brain. Nature biotechnology 42, 719–730. 10.1038/s41587-023-01798-5.

53. Windrem, M.S., Schanz, S.J., Guo, M., Tian, G.-F., Washco, V., Stanwood, N., Rasband, M., Roy, N.S., Nedergaard, M., Havton, L.A., et al. (2008). Neonatal Chimerization with Human Glial Progenitor Cells Can Both Remyelinate and Rescue the Otherwise Lethally Hypomyelinated Shiverer Mouse. Cell Stem Cell 2, 553–565. 10.1016/j.stem.2008.03.020.

54. Challis, R.C., Ravindra Kumar, S., Chan, K.Y., Challis, C., Beadle, K., Jang, M.J., Kim, H.M., Rajendran, P.S., Tompkins, J.D., Shivkumar, K., et al. (2019). Systemic AAV vectors for widespread and targeted gene delivery in rodents. Nat Protoc 14, 379–414. 10.1038/s41596-018-0097-3.

55. Plog, B.A., Mestre, H., Olveda, G.E., Sweeney, A.M., Kenney, H.M., Cove, A., Dholakia, K.Y., Tithof, J., Nevins, T.D., Lundgaard, I., et al. (2018). Transcranial optical imaging reveals a pathway for optimizing the delivery of immunotherapeutics to the brain. JCI Insight 3. 10.1172/jci.insight.120922.

